# Optoretinography in R9AP-bradyopsia reveals the essential role of G-protein signaling in the human cone elongation response

**DOI:** 10.64898/2026.04.28.721497

**Authors:** Connor Weiss, Teng Liu, Vimal P. Pandiyan, Yan Cao, Debarshi Mustafi, Joseph Carroll, Kirill Martemyanov, Edward N. Pugh, Ramkumar Sabesan

**Author notes:** Co-corresponding authors:* Ramkumar Sabesan, University of Washington, 750 Republican St., E213, Seattle, WA 98109, Edward N. Pugh, Jr., University of California, Davis, 3301 Tupper Hall, Davis, CA 95616. equal contributions (Connor Weiss, Teng Liu).

## Abstract

Human cone and rod outer segments exhibit a rapid shrinkage followed by a slower elongation in response to light, forming the basis of the optoretinogram. The molecular basis of this optical assay of photoreceptor function remains incompletely understood. In mouse rods, the elongation response requires transducin, the G-protein α-subunit activated in the initial amplifying step of the phototransduction cascade. Here, we measured human cone outer segment responses in subjects with bradyopsia arising from a triple-deletion mutation in R9AP, the anchor and transport protein for the GTPase-activating protein (GAP) RGS9. Immunoprecipitation showed that the mutant R9AP has greatly reduced affinity for RGS9, predictably reducing the level of RGS9 in the outer segment. The bradyopsia subjects’ elongation responses had normal activation kinetics, amplitude and photosensitivity, but markedly slowed recovery. These results indicate that normal levels of RGS9 are required for deactivation of the cone elongation response, situating the underlying molecular mechanism within the G-protein cascade at or prior to formation of the GAP complex. The recovery of the cone elongation response, measured in a paired-flash paradigm for stimuli isomerizing up to 75% of opsin, revealed that elongation requires a substrate that is depleted and recovers in a bleach-dependent manner. In particular, recovery from the highest bleaching exposure (75%) tracked the time course of cone opsin regeneration, implying that unregenerated cone opsin produces “*dark light*”, known in rods to arise from constitutive activation of G-protein. Taken together, the results from controls and bradyopsia identify the inactive (GDP-bound) transducin complex as the essential substrate for the human cone elongation response. Suppression of the elongation response in the paired-flash paradigm further revealed the complete profile of the initial fast cone outer segment shrinkage response and reinforced its origin in opsin structural changes. Overall, these results identify key mechanistic elements of the optoretinogram and enable its use as a molecularly interpretable, non-invasive assay of cone function in disease and therapeutic response.

## INTRODUCTION

Cone and rod photoreceptor outer segments undergo light-dependent changes in length that can be observed with nanometer scale resolution by phase-resolved optical coherence tomography (OCT) (Hillmann et al. 2016; Azimipour et al. 2020; Boyle et al. 2020; Pandiyan et al. 2020; Ma et al. 2021; Cooper, Brainard, and Morgan 2020; Zhang et al. 2019; Pandiyan, Nguyen, et al. 2022b; Zhang et al. 2017; Tomczewski et al. 2025). These outer segment length changes and correlative changes in scattering at the base and tips of the outer segments are now known as the optoretinogram (ORG), in analogy with the electroretinogram (ERG), the field potential generated by the combined electrical responses of rod and cone photoreceptors and downstream retinal cells (Pugh 1998; Robson et al. 2003). The human cone ORG can be decomposed into three components, distinguishable by their sign (decrease vs. increase in outer segment length), kinetics and photosensitivity (Pandiyan, Nguyen, et al. 2022b; Boyle et al. 2020; Pandiyan et al. 2020). The Shrinkage Response (Component 0), an initial fast reduction in cone outer segment optical path length (OPL), has been hypothesized to arise from lamellar disc deformation consequent to the internal charge displacement in cone opsins triggered by chromophore isomerization (Boyle et al. 2020; Pandiyan et al. 2020). The Slow Elongation (Component 2) is also stoichiometrically related to opsin bleaching, and has been hypothesized to arise from water entry into photoisomerized opsins driven by conformational changes that generate sub-molecular “pockets” into which water molecules diffuse (Pandiyan, Nguyen, et al. 2022b; Chawla et al. 2021).

The rapid Elongation Response (Component 1), an amplified lengthening of the outer segment, has been hypothesized to arise from water osmotically driven into the outer segment by free phosphate (P_i_) produced by the regulator of G protein signaling 9 (RGS9)-catalyzed hydrolysis of GTP in the activated G_tc_α-GTP-PDE6-RGS9 complex. To investigate this hypothesis, we measured ORGs of subjects with bradyopsia arising from an in-frame triple deletion in *R9AP*, the anchor protein for the RGS9-Gβ5L GTPase Activating Protein (GAP) complex (Michaelides et al. 2010; Nishiguchi et al. 2004). As binding of RGS9-Gβ5L to R9AP is necessary for transport of the GAP complex to outer segments, defects in R9AP are predicted to slow the rate of GTP hydrolysis (Hu and Wensel 2004; Hu and Wensel 2002; Martemyanov et al. 2003; Keresztes et al. 2004; Burns and Pugh 2009; Krispel et al. 2006) and thereby reduce the rate and quantity of P_i_ production. We tested this prediction by measuring the activation and recovery kinetics of the elongation response with stimuli producing 8 to 75% pigment bleach for up to 15 s in a single-flash protocol, and over periods up to 2 min with a novel paired-flash protocol. Together, the results reveal previously unresolved dynamics of cone outer segment shrinkage and elongation that accompany its light response. Comparison of the ORGs of R9AP-bradyopsia subjects with controls further provides new constraints on the biophysical and biochemical mechanisms underlying the human cone ORG.

## RESULTS

### The recovery of the human cone elongation response is slowed in R9AP-bradyopsia

The bradyopsia subjects of the study are siblings with a homozygous in-frame deletion in the *R9AP* gene (c.94_102del, p.Asp32_Gln34del) (Fig. 1)(Michaelides et al. 2010). Their cone mosaic in the fovea and near temporal periphery were remarkably normal in appearance (Fig. 1A), with cone density within the normal range (Fig. 1B). In the retinal region from which ORGs were measured, the average lengths of the probands’ outer segments were 23.3 and 22.0 μm respectively, not reliably different from the normal average, 21.5 μm (Table S1). Cone density remained stable over the five years between the current and a previous (Strauss et al. 2015) investigation.

**Figure 1.**
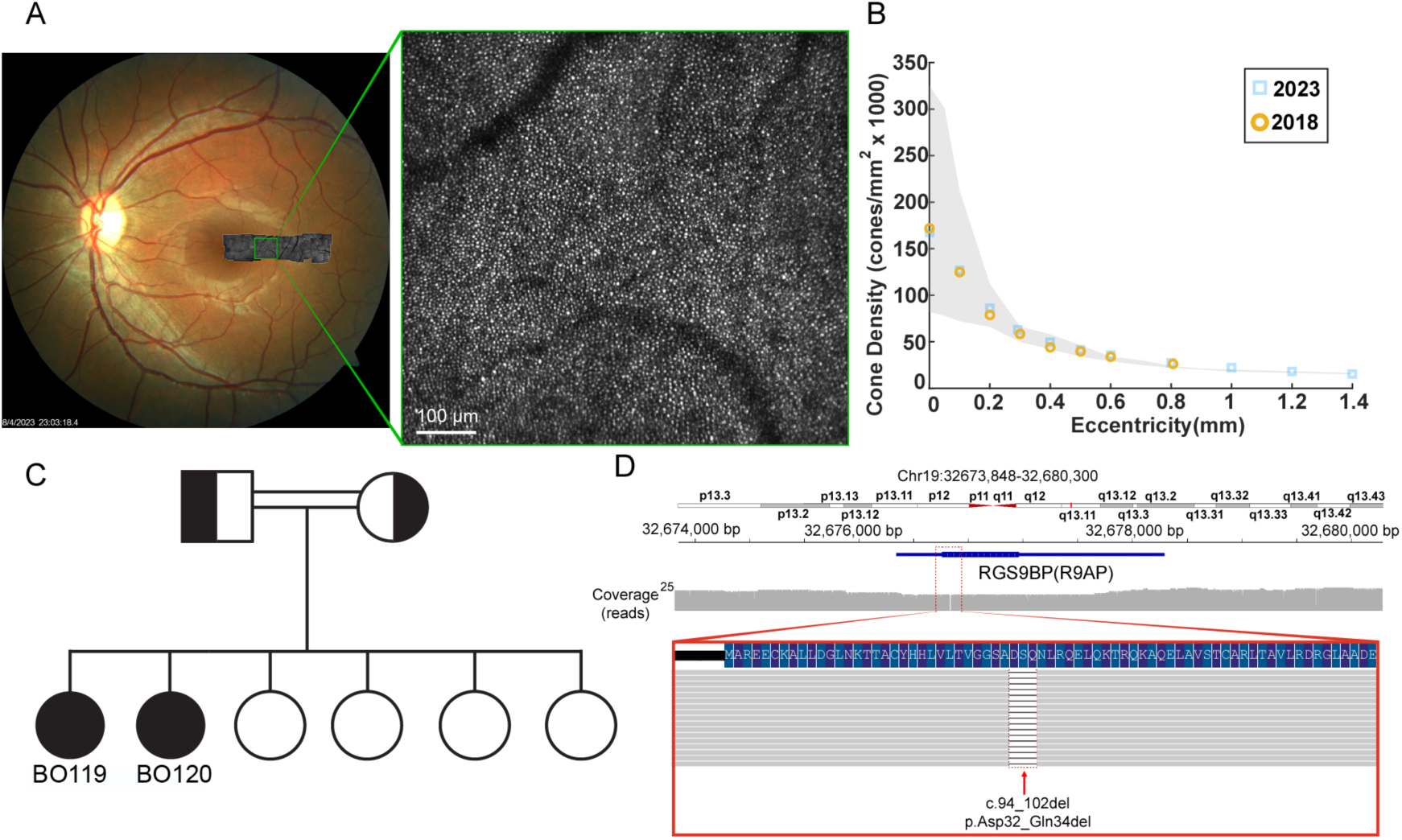
Genetic and phenotypic characterization of the R9AP-bradyopsia patients of the study. **A**. Fundus image of bradyopsia subject BO119, showing the temporal eccentricity range where AOSLO confocal imaging was performed. The green-highlighted square is magnified to show the cone mosaic. **B.** Cone density of Subject BO119 at varying eccentricity measured in the year 2018 (orange) and in 2023 (blue). The range of normal cone density measured with histology(Curcio et al. 1990) is indicated in the gray shaded area. **C.** Pedigree of the bradyopsia subjects, BO119 and BO120. **D**. Long-read sequencing highlights the homozygous pathogenic in-frame deletion in *R9AP* in subjects BO119 and BO120 (cf MATERIALS AND METHODS, *Long-read genome sequencing*).

Cone ORGs of the subjects with R9AP-bradyopsia and those of normal controls have comparable activation kinetics, amplitudes and photosensitivities (Fig. S1, S2, Table S1), revealing that these bradyopsia subjects’ cones are functional. However, the recoveries of their ORG responses to stimuli that bleach small fractions of cone pigment are substantially slowed relative to those of normal controls (Fig. 2A). The half-times of recovery of two normal subjects to stimuli delivering 1.8×10^6^ photons μm^-2^ are 3.2 s and 2.9 s (filled green and red circles in Fig. 2A, bottom), whereas the recovery halftime of bradyopsia subject BO119 to a flash of 1.9×10^6^ photons μm^-2^ is 8.2 s (Fig. 2A, top, blue open circle). Similarly, the recovery halftime for a control subject to a stimulus delivering 3.2×10^6^ photons μm^-2^ is 6.8 s (Fig. 2A, bottom, filled green square), whereas the recovery halftime for a bradyopsia subject (BO120) to 3.5×10^6^ photons μm^-2^ is 11.1 s (Fig. 2A, open orange square). An even more telling comparison of the response recoveries of bradyopsia subject BO119 and controls is provided by overlaying the traces (Fig. 2B): the bradyopsia and control ORGs rise on the same trajectories, but the control traces diverge from the common rising phase at ∼0.7 s with initial slopes of −40 nm s^-1^ (red trace) and −25 nm s^-1^ (green traces), while the bradyopsia subjects’ recoveries have a 2- to 3-fold lower magnitude slope of −13 nm s^-1^ (blue data points and cyan lines).

**Figure 2.**
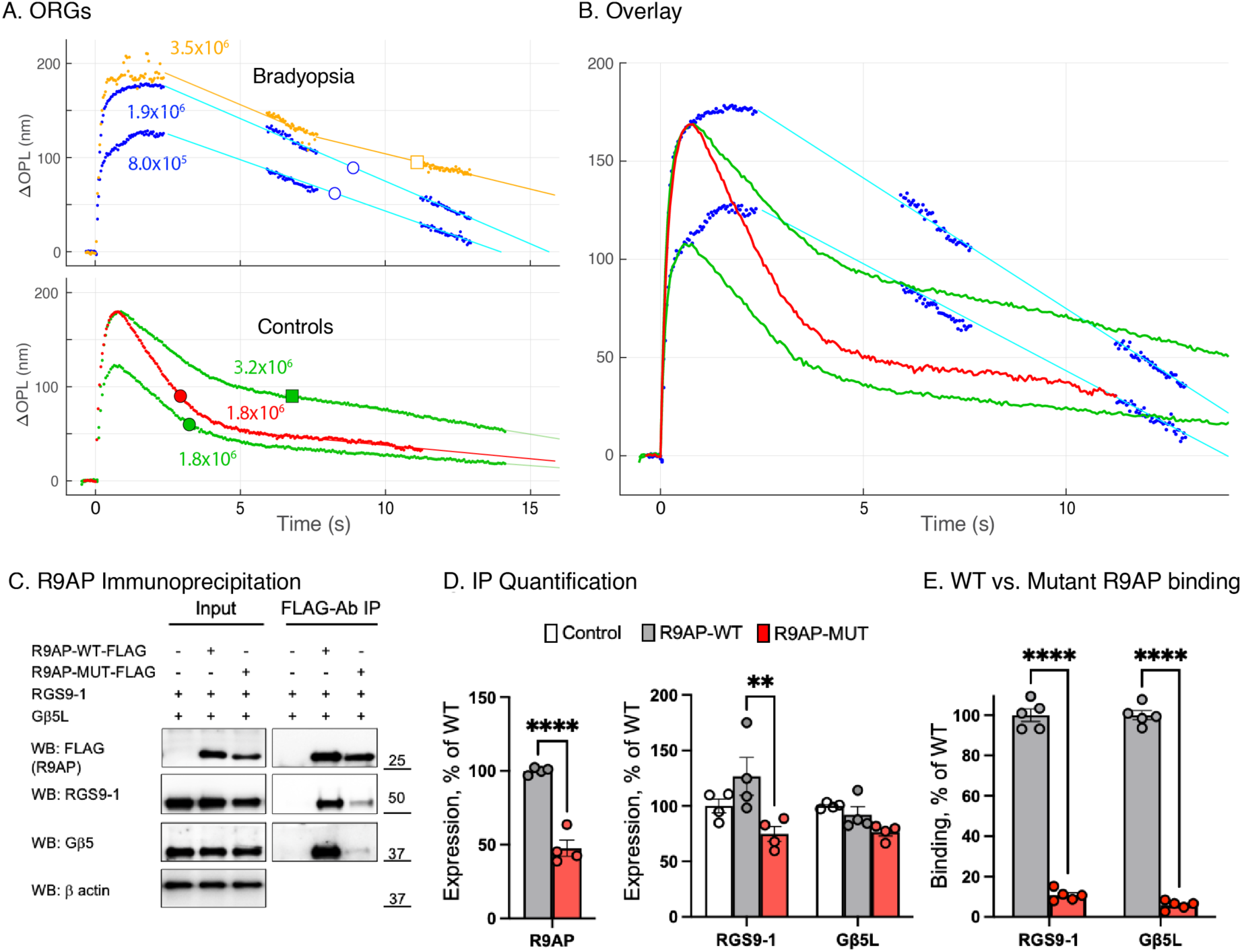
Slowed recovery of the cone Elongation Response of subjects with R9AP-bradyopsia is attributable to reduced RGS9-1-Gβ5L in the patient’s cone outer segments. **A.** *Upper plot*: ORGs of two subjects (BO119, blue symbols; BO120, orange) with the R9AP deletion mutation (Fig. 1); stimulus energy densities are adjacent to the traces. Symbols are plotted at the times of 50% recovery, and straight lines have been fitted to extract slopes (gaps from brief, planned respites). *Lower plot*. ORGs of two control subjects, with symbols plotted at the recovery halftimes. **B**. Overlay of the ORGs of subject BO119 replotted from panel A (blue dots) and those of the two control subjects (replotted as uninterrupted lines; green traces were scaled by 0.90 and 0.94, respectively and red trace by 0.94, which brought the rising phases into correspondence). **C – E**. Experiments in HEK293T cells reveal that ΔAsp32-Gln34 R9AP is translated, but the protein has a reduced capacity to bind the RGS9- 1/G5βL complex. **C**. Co-expression and co-IP of R9AP with the RGS9-1/G5βL complex. HEK293T cells were transfected with the specified constructs and lysed 24 hours post-transfection in a buffer containing 1% Triton X-100. IP was performed with anti-Flag affinity resin, and the eluted proteins analyzed by Western blotting with specific antibodies. (Cells transfected with all constructs except the Flag-tagged R9AP were used as controls to assess non-specific binding.) **D**. Quantification of protein in lysates of cells transfected with the specified constructs. Protein quantities normalized relative to β-actin are shown as mean ± SEM (n=4): **, *p*<0.01, two-way ANOVA followed by Sidak’s multiple-comparisons test (RGS9-1 and Gb5L); ****, *p* < 0.0001, un-paired *t*- test (R9AP). **E.** Quantification of the amounts of RGS9-1 or Gβ5L bound to R9AP. Data (mean ± SEM, n=5) normalized to amount of R9AP in eluate; 2-way ANOVA followed by Sidak’s multiple-comparisons test: ****, *p* < 0.0001.

R9AP is critical for normal function of the RGS9 GAP complex, since it anchors the complex to the disc membranes and is required for its transport to the outer segment (Martemyanov et al. 2003; Keresztes et al. 2004). We examined the impact of the 3 amino acid in-frame deletion mutation of the bradyopsia subjects on the expression of the mutant R9AP and on its interaction with the RGS9-1-Gβ5L complex with transfected HEK293 cells, investigating specifically whether the mutated gene is translated to a protein product that binds to the GAP complex (Fig. 2C, D). The mutant R9AP was found to be expressed at a lower level than the wild type (WT) protein (Fig. 2C, “input”), and immunoprecipitated much less RGS9-1 and Gβ5L than did WT-R9AP (Fig. 2C, “FLAG-Ab IP”). Quantification of the immunoprecipitation data revealed a substantial reduction of binding of RGS9-1 by the mutant R9AP (Fig. 2DE). These results support the conclusions that RGS9-1 and Gβ5L are reduced from the normal level in the bradyopsia subjects’ cone outer segments. Given that the R9AP mutation predictably reduces RGS9 levels in the outer segment, the earlier divergence of the control ORGs from the shared rising phase observed in the bradyopsia subject (Fig. 2B) is attributable to higher RGS9 levels in control COS. As RGS9 acts as the enzyme in a Michaelis complex in living photoreceptors (Burns and Pugh 2009; Krispel et al. 2006), this accelerated recovery is thus directly attributable to a higher RGS9-GTPase activity in the controls. Moreover, the linear initial phase of the recoveries of both normal and bradyopsia subjects suggests that the RGS9-GTPase reaction is substrate-saturated in both cases.

These results (Fig. 2) reject the hypothesis (Pandiyan, Nguyen, et al. 2022b) that free phosphate (P_i_) produced by RGS9-catalyzed GTP hydrolysis acts as an osmolyte to drive outer segment swelling. Thus, if osmotic swelling due to increase in P_i_ were responsible for the elongation response, the reduced RGS9 level in the bradyopsia subjects’ outer segments should have diminished the response’s initial velocity and amplitude but failed to do so (Fig. 2B; Fig. S1). Nonetheless, the bradyopsia ORGs reveal that the elongation response arises from one or more steps in phototransduction that precede RGS9- catalyzed GTP hydrolysis. This revelation led us to investigate the recovery kinetics of the elongation response over a wide range of bleaching exposures to identify additional features that would constrain its molecular mechanism.

### A paired-flash paradigm uncovers the time course of depletion and recovery of a substrate of the cone elongation response

Accurately measuring the ORG over an epoch of 10 s or longer (Fig. 2A, B) is challenging for several reasons, including eye blinks, maintaining fixation, and the difficulty of sustaining recordings of adequate bandwidth for long durations. To circumvent these limitations and further characterize the recovery of the elongation response, we developed a paired-flash protocol in which the ORG was recorded in response to an initial “Test” stimulus of one intensity followed at prescribed interstimulus intervals (ISIs) by a “Probe” stimulus of the same or a different intensity (Fig. 3A). Examples of the responses to the probe flashes for different subjects, ISIs and stimulus intensities are provided in Fig. 3C-H. To summarize the paired-flash ORG recovery results, we plotted the average amplitudes and initial velocities of the responses to the probe flash as a function of ISI (Fig. 3I, J). For all stimulus intensities, the recoveries of the ORG amplitude are well described by first-order exponentials, while the recoveries of the initial velocities follow two-phase trajectories, with an initial fast phase followed by an increasingly slower phase. The dependencies of the recovery time constants and initial rates on bleach level followed the general rule that the greater the bleach level, the slower the recovery (Fig. 3K, L). Specifically, the time constant of amplitude recovery increases with the proportion of pigment bleached (Fig. 3K), such that for a 75% bleach (red symbols), it approaches the time course of cone pigment regeneration (τ ∼ 2 min) (Hollins and Alpern 1973; Lamb and Pugh 2004).

**Figure 3.**
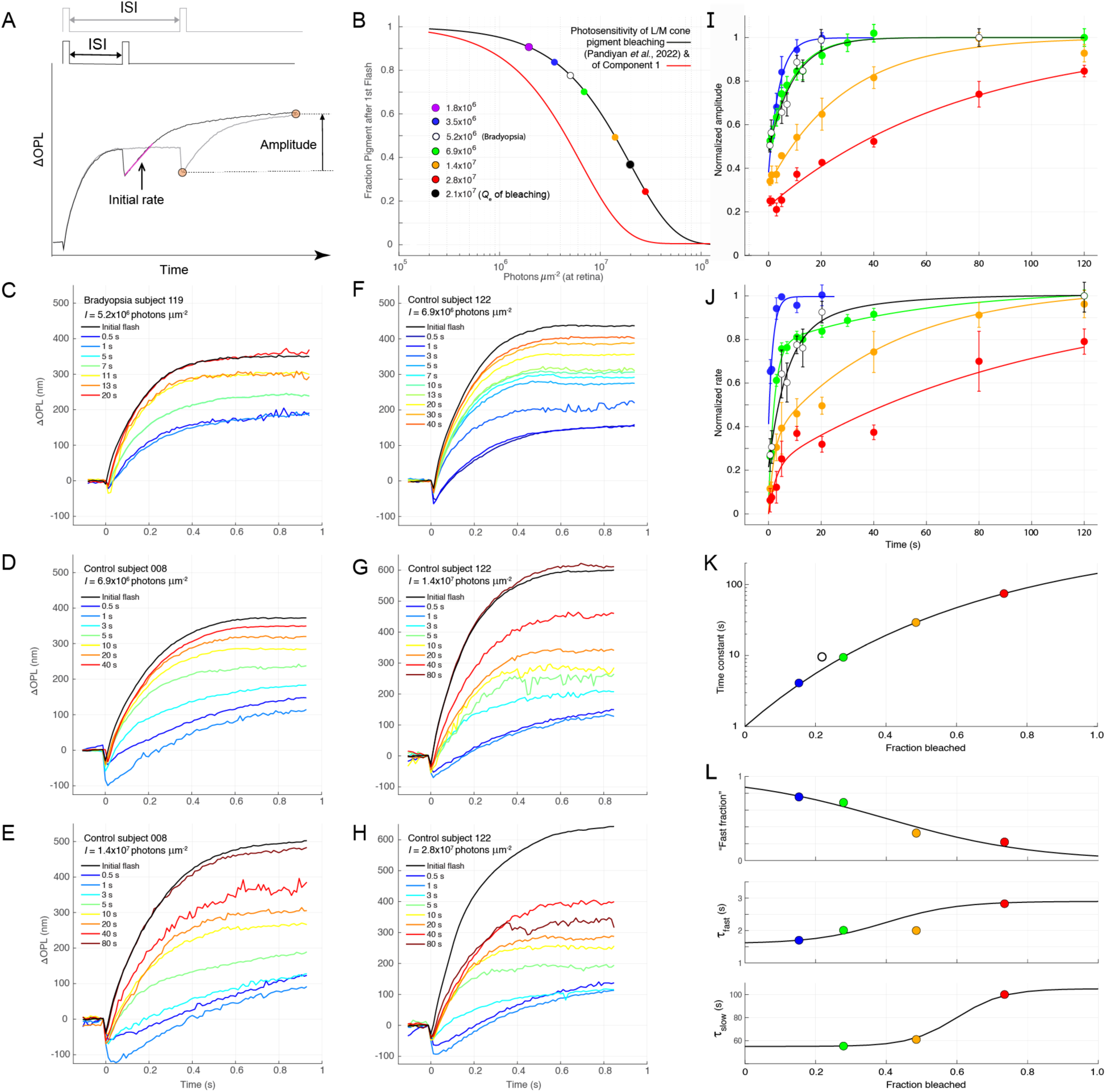
ORG paired-flash paradigm reveals that strongly bleaching stimuli deplete a substrate underlying the cone Elongation Response and allow the dependence of recovery on the fraction of cone opsin bleached to be characterized. **A**. Schematic of the paired-flash paradigm. In each trial, a Test flash of specific strength is initially delivered, followed by a Probe flash of the same flash strength, delivered at different interstimulus intervals (ISI). The symbols on the plot show how the amplitude of the probe response is measured, with the 1^st^ symbol indicating the baseline level, and the ΔOPL between the 1^st^ and 2^nd^ symbols taken to be the probe response amplitude. In addition, the initial slope (magenta line) of the response to the Probe is measured by least-squares fitting of a straight line to the initial portion of the response. **B.** Quantity of cone opsin bleached by the stimuli used in the paired-flash experiments. The smooth curve plots the function *P*(*Q*) = exp(–*Q*/*Q*_e_), where *P* is the fraction of cone pigment remaining after exposure to a stimulus of energy density *Q* photons μm^-2^, *Q*_e_ = 2×10^7^ photons μm^-2^ and 1/*Q*_e_ is the photosensitivity of cone pigment bleaching directly measured by Pandiyan *et al*. (Pandiyan, Nguyen, et al. 2022b) with adaptive optics serial-scanning, single-cone reflectometry (the fraction bleached is 1– *P*). The symbols plot the fraction of pigment remaining for the different energy densities used in the paired-flash experiment and correspond in color to the symbols in panels J – L. The red curve plots the complement of the exponential saturation function that describes the normalized amplitude vs. intensity function describing the Elongation Response of all the subjects of the study (Fig. S1); shifted left from the bleaching curve by a factor of 5, it represents the fraction of a hypothetical substrate that remains unactivated after a bleaching exposure specified by the abscissa. **C – H.** Families of responses to the 2^nd^ (Probe) flash for different subjects and different flash strengths, identified on the plots. **I.** Plots of the recovery of the amplitude (see panel A) of the ORG response to the 2^nd^ flash from the experiments of C-H (and others with different subjects) for 4 different flash strengths: the amplitude of the response to the 2^nd^ flash was normalized by the initial, dark adapted amplitude *(cf* black traces in C-H); each curve presents the mean ± std. deviation for 3 replications with different subjects, except for the lowest intensity (blue symbols, n = 2). The amplitude recovery data for the different flash strengths were fitted with single rising exponential curves whose time constants are plotted in K. **J.** Recovery of the initial velocity (rate) of the responses to the 2^nd^ flash. For all stimulus energy densities the recovery is characterized by two phases, with the first phase much faster than the second (curves). A “slowdown” phenotype of the bradyopsia patient’s ORG is seen by comparing the recovery of the initial velocity with that of controls to a 30% more intense stimulus: the bradyopsia patient’s recovery is prolonged by ∼ 5 s during the initial 10 s after the initial stimulus. **K.** Time constants of the recovery of amplitude in panel I for different stimulus strengths, plotted as function of the fraction of pigment bleached (panel B). **L**. Parameters of the curves describing the recovery of the initial velocity of the ORGs in J plotted as function of the fraction of pigment bleached. The two lower subpanels plot the fast and slow time constants of recovery, while the upper subpanel plots the “fast fraction”, the fraction of the recovery governed by the fast time constant.

The paired-flash data, taken together with the slowed recovery of the bradyopsia subjects (Fig. 2A, B), reveal that a depletable substrate (Fig. 3B, red curve) required for the RGS9 reaction is produced at the same initial rate and total quantity for each flash strength in bradyopsia and control subjects, and that *recovery* represents the restoration of the dark-adapted level of this substrate. One candidate for such a depletable substrate of phototransduction that precedes the RGS9-catalyzed GTP hydrolysis is cone G_tc_, the inactive (GDP-bound) heterotrimeric G-protein itself. The ratio of G_t_α in the rod outer segment has been measured to be 1:8 relative to rhodopsin (Lobanova et al. 2008), and that of G_tc_:P in cones is assumed to be similar. That the time course of amplitude recovery of the elongation response after a 75% bleaching stimulus appears to track cone pigment regeneration suggests that one or more species of bleached pigment depletes this substrate, causing *dark light*, i.e., persistent G_tc_ activation that is ultimately eliminated only by regeneration of the opsin itself, known to have a time constant of ∼ 2 min (Hollins and Alpern 1973; Lamb and Pugh 2004; Yao, Fay, and Farrens 2025).

### Testing the hypothesis that the elongation response corresponds to an activated state of cone G-protein

The hypothesis that the elongation response is instantaneously proportional to an activated species of cone G-protein is suggested by its properties: (1) a saturating response vs. intensity function that is linear for low flash strengths, but with photosensitivity 3-fold greater than that of bleaching (Fig. 3B, red curve; Fig. S1; (Pandiyan, Nguyen, et al. 2022b); (2) rapid, activation kinetics (τ = 175 ms; Fig. S1, Table S1); (3) recovery kinetics slowed by R9AP-bradyopsia that reduces RGS9 in the cone outer segment (Fig. 2; Fig. S2). To investigate this hypothesis, we used the paired-flash protocol to characterize the “dim-flash” elongation response, i.e. its activation and deactivation kinetics for a flash whose intensity (1.8×10^6^ photons μm^-2^, Fig. 3B) is in the linear regime of response vs. stimulus intensity function (Fig. 4; Fig. S1).

**Figure 4.**
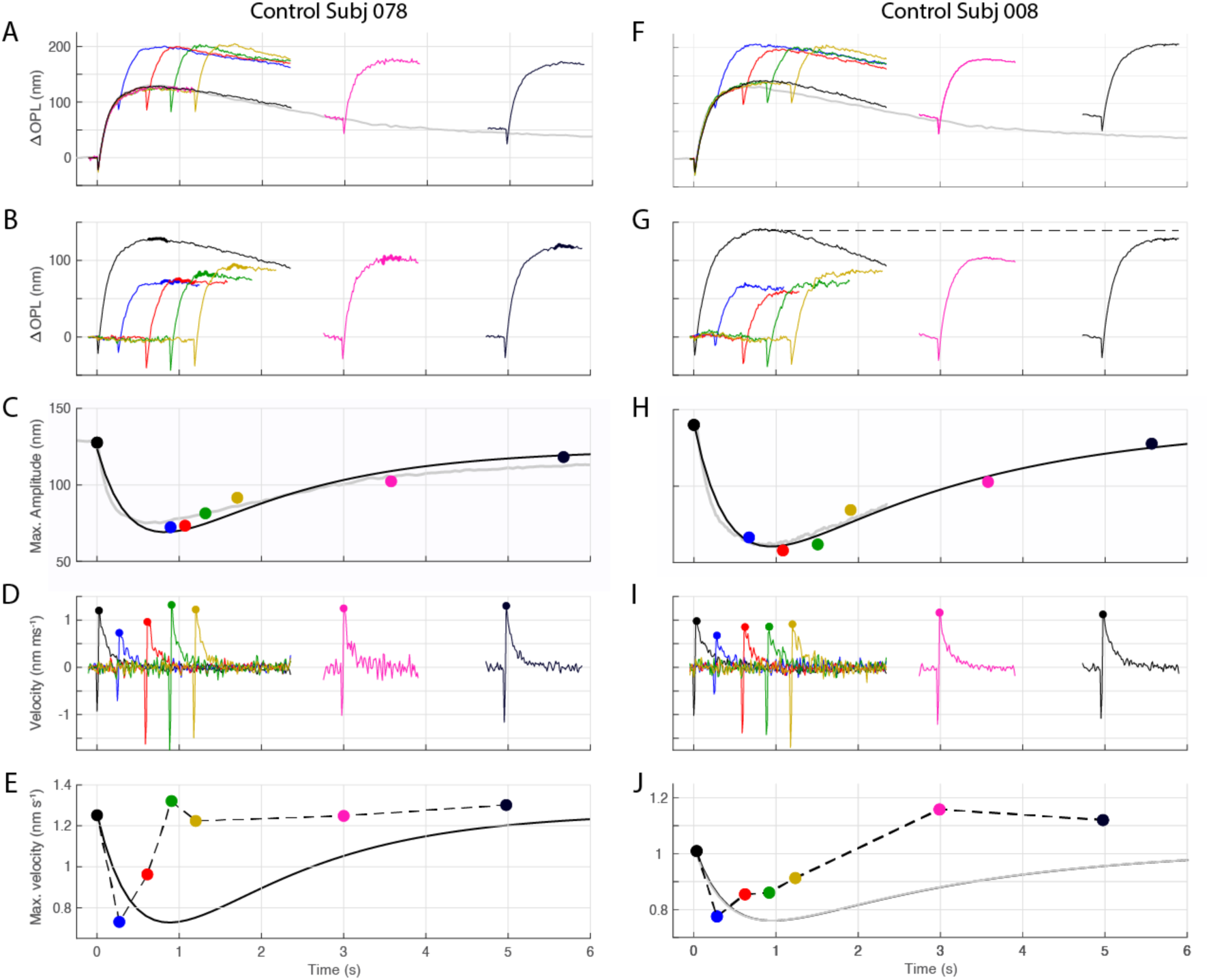
Paired-flash measurements of the Elongation Response recovery in a experiment with a stimulus (1.8×10^6^ photons μm^-2^) that bleaches 8% of the cone pigment (Fig 3B). **A, F.** Results for two subjects. Each differently colored trace presents the average of ORGs measured in trials (n = 4–5) that delivered paired flashes initially at *t* = 0 and at again at the various times when the trace again rises. The black trace is the test flash response for each subject (n = 10, and 9 trial average respectively), and extends uninterruptedly to ∼ 2.2 s. The gray trace in A and F is replotted from Fig. 2B, and was obtained in separate experiments with the same flash strength from the subject of panel A. The dashed lines provide fiducials for the maximum amplitude of the test flash response in subsequent plots of the same column. **B, G.** Incremental ORG responses extracted from the traces in A, F by subtraction of the test flash response from the trace for ISIs < 2 s (blue, red, green, gold, pink for ISIs ranging from 0.3 - 3 s, the responses to the second flash were shifted by baseline subtraction to the zero level, and their maximum amplitudes measured as the average over bands (thickened portion of traces) containing the apparent maxima.) The horizontal dashed lines are repeated from A, F to provide fiducials of the subjects’ ORG maximum response in the dark-adapted state. **C, H.** Maximum amplitudes from B, G plotted a function of ISI (the symbols are plotted at the midpoint of the region from which they were extracted). The gray curve in C replots the gray trace in A, inverted, offset to the fiducial line (129 nm) and scaled by 0.44; the gray curve in H replots the control response in F, inverted, offset to the fiducial (141 nm) and scaled by 0.55. The smooth black curves plot a bi-exponential curve, *A*(*t*) = *A*_o_ {1 – [exp(–*t/*τ_1_ – exp(–*t/*τ_2_)]} fitted to the data by least-squares with the Matlab^TM^ “cftool” GUI: for the curve in C, *A*_o_ = 123 nm, τ_1_ = 0.47 s and τ_2_ = 1.67 s; for that in H, *A*_o_ = 141 nm, τ_1_ = 0.44 s and τ_2_ = 2.54 s. **D, I**. Derivatives of the ORG traces in B, G calculated with the Matlab^TM^ “gradient” function; the symbols plot the maxima of the derivative curves. **E, J.** The maxima of the derivatives in D, I are plotted at the times of the maxima. The dashed curve simply connects the points while the black curves are repeated from C, H, but scaled to match the minimum velocity (blue circles), showing that the initial velocity of Elongation Response recovers more quickly than the amplitude.

The decreases in amplitude of the elongation response caused by an initial stimulus and extracted from the probe flash responses (symbols in Fig. 4C, H) are well approximated by an inverted, scaled version of the response to the initial test flash (gray traces in C, H). A priori, there is no reason to expect the response to obey this scaling relation, but its effectiveness in describing the probe-response amplitude decrements suggests that not all forms of activated G-protein contribute to the elongation response. Thus, if the amplitude of the response, *C*_1_(*t*), is directly proportional to the total quantity *G*\*(*t*) of activated G-protein, then at time *t* during response to the initial flash

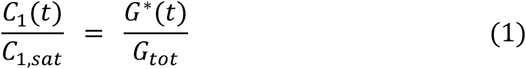

where *G*_tot_ is the total quantity of G-protein and *C*_1,*sat*_ is the saturated elongation response magnitude. If the protein is present in only two forms, inactive (*G*_0_, cone G_tc_-GDP) and active, then at the time *t*_ISI_ of a probe flash, the quantity of inactive protein is

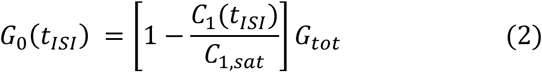

If P* produced by a probe flash generates *G*\*(*t*) at a rate proportional to the quantity of inactivated protein, then the amplitude of each probe response will be proportional to *G*_0_(*t*_ISI_) (Eq 2), and the decremented amplitudes of the probe flash response are thereby predicted to track a scaled form of the original response. The quantity in the square bracket of Eq 2 predicts not only the kinetics, but also the absolute magnitude of the scaling: at the time of the probe response minimum (*t*_ISI_ ∼ 0.8 s) the scale factor (square bracket in Eq 2) is predicted to be 0.75 and 0.76 respectively for the two subjects, but the scaling factors used for the fitting were 0.44 and 0.55 (Fig. 4C and H, respectively). This scaling mismatch is interpreted to mean that in addition to the form of activated G-protein that is proportional to the elongation response for this stimulus, there is at least one other form comprising up to 40% of the total activated quantity that does not contribute to the elongation response to the probe. This conclusion adjusts upward the rate of activation per P* that would be estimated from assuming the elongation response is directly proportional to the activated state in a 2-state (inactive/active) model of G_tc_.

The dim-flash recovery of the elongation response is well described as the convolution of two first-order exponential decays (black curves in Fig. 4C, H), with “fast” time constants of 0.47 s and 0.44 s, and “slow” time constants of 1.67 s and 2.54 s for the two subjects, respectively. In models of phototransduction, the activation and deactivation of the G*-E* (G_tc_α-PDE) complex that produces light-activated cGMP hydrolysis obeys similar bi-exponential kinetics (Gross, Pugh, and Burns 2012; van Hateren and Lamb 2006), with one time constant assigned to deactivation of photoactivated opsin and the other to deactivation of G*-E*, but a priori assignment of the two time constants to the underlying biochemical reactions is not possible due to the formal interchangeability of the time constants. However, given identification of the depletable substrate of the elongation response as the inactive, heterotrimeric cone transducin, and the bi-exponential deactivation of the dim-flash elongation response, it can be concluded that the complete cone G-protein cycle - from activation by P*, to activation of PDE, RGS9-catalyzed GTP hydrolysis and reconstitution of the transducin complex - occurs in no more than 2.5 s.

### Characterization of the cone Shrinkage Response with the Paired-Flash Paradigm

In single-flash experiments, the initial, negative-going shrinkage response (Component 0) is truncated by the larger and more photosensitive positive-going elongation response. Suppression of the latter in the paired-flash paradigm can unmask the shrinkage response. This unmasking effect is evident in the first few seconds after an initial stimulus, where the magnitude of the shrinkage response to a second flash of the same intensity delivered at ISI = 1.2 s was ∼3-fold greater than its initial, dark-adapted magnitude (Fig. 3D –H, blue traces). We exploited this effect to reveal the complete form of the shrinkage response with a paired-flash protocol where the intensity of the initial test flash was chosen to largely suppress the elongation at the time of delivery of a second probe flash of varied strength (Fig. 5A-F).

**Figure 5.**
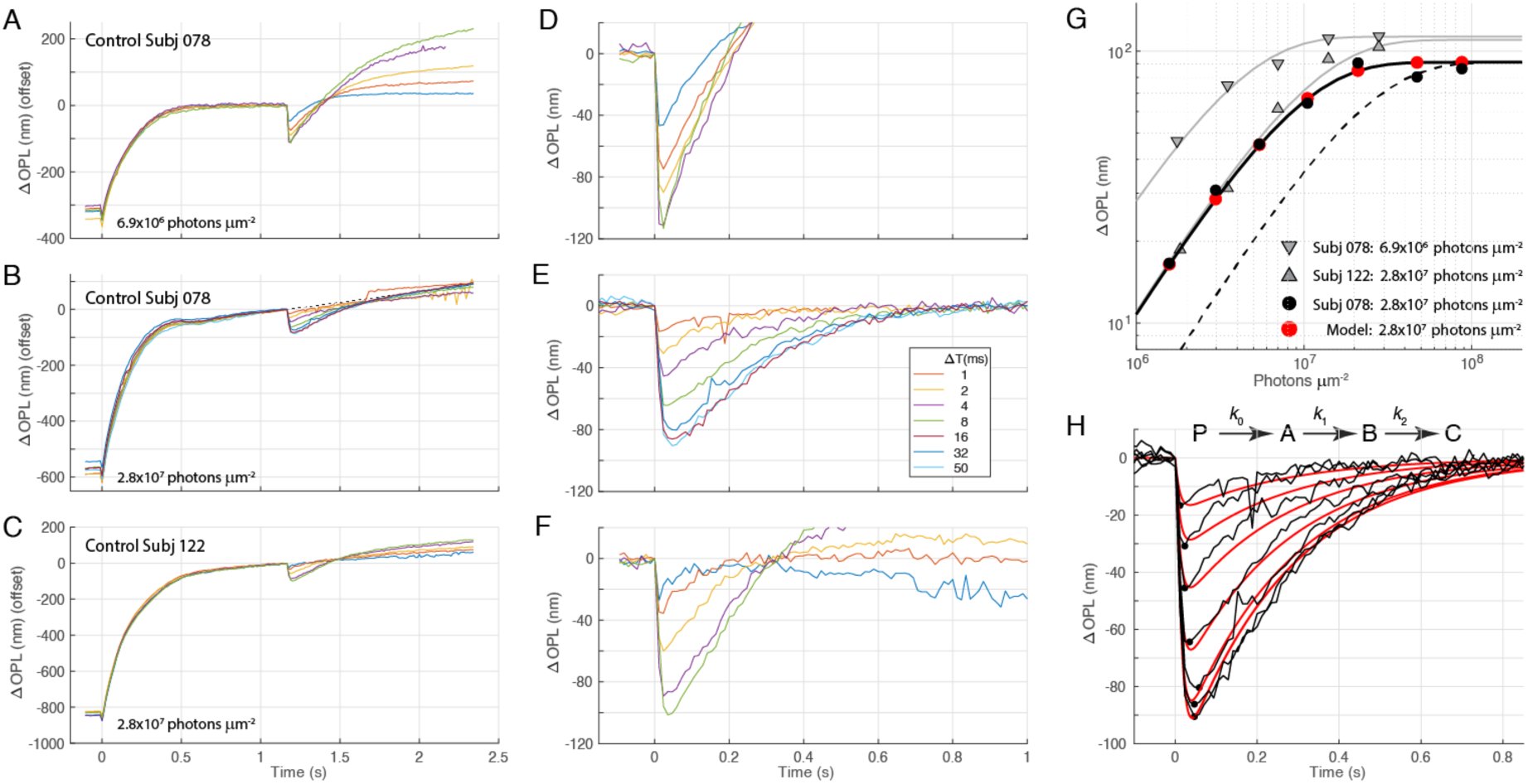
Kinetics, amplitude and photosensitivity of negative-going ORG Component 0 obtained from a paired-flash paradigm. **A– C**. ORGs in response to an initial test flash of fixed intensity (indicated in the lower left of each panel) and a variable-strength probe flash delivered at 1.2 s. All traces have been offset so that they are at zero level at the time when the second flash was delivered. **D – F.** ORG families obtained from the responses to the probe flashes extracted from the corresponding panels to the left and presented on expanded abscissa and ordinate axes. For the data in panel B following the probe flash, all the traces relaxed to a common line (dashed) that was a continuation of the trace preceding the probe flash; the traces in E were obtained by subtracting this common line from them. For the ORGs in A and C, a common baseline was not present. **G.** Peak magnitudes (absolute values) of the Component 0 traces in panels D – F; the gray triangles plot the peak magnitude of the data in panels A and C, respectively, while the filled black symbols plot the magnitudes of Component 0 responses in panel E. The smooth curves fitted to the Component 0 magnitudes are exponential saturation functions *M*(Q) = *M*_max_ [1 – exp(–*Q*/*Q*_c_)], with log_10_(*Q*_c_) = 6.54 (gray up triangles), 7.01 (gray down triangles), 6.9 (filled black symbols), and 7.3 (dashed line, corresponding to the bleaching function in Fig. 3B). **H**. Kinetic scheme and predictions (red curves) of the Component 0 results from panel E (replotted as black traces). The kinetic scheme embodies the hypothesis that isomerization causes the cone opsin to transition sequentially through distinct states, beginning with *P*, the pigment present when the bleaching stimulus is delivered. The intermediate state *B* is identified as corresponding to Component 0. The red curves represent solutions for the rate equations describing the kinetic scheme for bleaching exposures of the same power density (1.75×10^9^ photons μm^−2^ s^−1^) but with varied durations of 1, 2, 4, 8, 16, 32 and 50 ms. Panels AD and BCE are obtained with stimuli of the same power density with durations equal to 4 and 16 ms respectively. A notable feature of the ORG data captured by a model is that for the three longest duration stimuli the magnitudes are nearly identical, i.e., they are saturated (cf. also panel G, filled symbols), as expected if the cone pigment is fully bleached by these stimuli.

The utility of maximally suppressing the cone elongation response to characterize the shrinkage is highlighted by comparison of results obtained from one subject for initial stimuli differing by a factor of 4 (Fig. 5D vs. 5E): the 2^nd^-flash ORGs turn upward much more rapidly after the lower intensity initial stimulus (Fig. 5D) than after the more intense first flash (Fig. 5E). We hypothesize that truncation of shrinkage response in 5D for the second probe flash is caused by activation of the remaining substrate underlying the elongation response not yet activated by the first test flash. This hypothesis is supported by a comparison of the elongation amplitudes for the initial test flash: elongation has an initial amplitude of ∼ 310 nm in Fig. 5A, but the amplitude is ∼500 nm in Fig. 5B, essentially in saturation (Fig. S1). Thus, we conclude that in the experiment of Fig. 5A, D, there is substantially more elongation response substrate available to be activated by the probe stimulus than in the experiment of Fig. 5B, E.

A second line of support for this conclusion is the much higher velocity of the positive-going, truncating portion of the ORGs in 5D than in 5E. Complete suppression of elongation response by the initial flash uncovered a Component 0 response family to second flashes of varied strength whose features link it to opsin changes after bleaching. First, the Component 0 responses to the 3 strongest stimuli had essentially the same kinetics and magnitude, despite a 3-fold range in flash duration (16 to 50 ms): this saturation behavior is expected if the three stimuli bleach essentially all the available cone visual pigment by ∼ 16 ms. Second, a kinetic model based on the general principle that photoisomerized cone pigment transitions through intermediate structural stages well describes the Component 0 response family (Fig. 5H red traces). Because variation in the strength of the second flashes (Figs. 5D, E, F) was effected by variation in the stimulus duration, a necessary feature of the kinetic model is that the input has the form of a rising exponential, i.e., *P*(*t*) = *P*_0_ [1 – exp(−*k*_0_ *t*)], *t* ≤ Δ*T*, where Δ*T* is the stimulus duration, *k*_0_ = *I*_0_/*Q*_e_, where *I*_0_ is the power density of the stimulus (photons μm^-2^ s^-1^) and 1/*Q*_e_ is the photosensitivity (*cf* SI “Multi-state model of cone opsin bleaching and decay applied to Component 0 and Component 2 of the ORG”).

The saturated magnitude of Component 0 in Fig. 5D is 110 nm, surprisingly greater than that in Fig. 5E (85 nm), despite the rapid truncation of Component 0 by the elongation response in Fig 5D and the 4-fold stronger stimulus used in the experiment of Fig 5E. In the context of the hypothesis that the magnitude of Component 0 is proportional to an opsin decay intermediate, the lower magnitude of Component 0 with a higher strength stimulus of Fig 5D, E is explained by the observation that the initial stimulus bleached a substantial percent (∼75%: cf. Fig 3B, red symbol) of the cone pigment, so that even a fully bleaching second stimulus produced a relatively small fractional bleach, i.e., only 25% of the total dark adapted pigment. An initially puzzling feature of these results is that the photosensitivity (Fig. 5G, solid black symbols and curve) is leftward-shifted 2.5-fold from the previously measured photosensitivity of bleaching at this retinal locus (dashed curve)(Pandiyan, Nguyen, et al. 2022b). We propose that the resolution of this puzzle is that a strongly bleaching initial stimulus causes an increase in photosensitivity to a subsequent bleaching exposure: such an increase is expected based on the decrease in cone pigment axial density accompanying substantial bleaching (Fig S4; SI, “Increase in the photosensitivity of bleaching by strong prior bleaches”; Pandiyan *et al*., 2022, Fig. S3 in the reference), but could involve additional mechanisms.

In summary, several challenges arise in the empirical characterization of the outer segment shrinkage Component 0, and in linking it to photopigment bleaching, including the inescapable truncation by the elongation response in the single-flash paradigm (e.g., Fig. 5D), the photosensitivity shift when elongation response has been fully suppressed by a prior stimulus that bleaches substantial pigment (Fig. 5G), and systematic differences between subjects in the magnitude of elongation response. An aspect of this latter variation is illustrated by the results of the two subjects: even after the delivery of a strong elongation response-suppressing stimulus, the positive-going relaxation of Component 0 from its minimum can display an “isosbestic” point, crossing from negative to positive at ∼ 300 ms after the stimulus (Fig. 5C, F). Such behavior cannot be accommodated by the subtraction of a constant sloping baseline as in Fig. 5B, E to retrieve the complete bleach dependence of Component 0. Rather, it suggests that the relatively rapid increase in ΔOPL immediately following the initial decrease may manifest a distinct state of the opsin that follows ineluctably from the state underlying the initial shrinkage, identified as Component 0.

### ORG Component 0 and Component 2 can be understood as successive states of photoactivated cone opsin

Responses to flashes delivered at ISIs of 0.5 s and 1.0 s in paired-flash experiments follow very similar trajectories (Fig. 3D-H, dark- and light-blue traces). For stimuli that bleach less than ∼ 20% of the pigment (cf. Fig. 3B), the responses at these early ISIs correspond to the elongation not activated by the initial flash. This follows, because the reciprocal photosensitivity of elongation response for all subjects is∼6.6×10^6^ photons μm^-2^ (Fig. S1), so that less than 100 × [1 − exp(−3×10^6^/6.6×10^6^)] = 36% of elongation response is activated by initial flashes producing less than 3×10^6^ photons μm^-2^. Consequently, a 2^nd^ identical flash at ISI = 1 s is expected to generate an ORG with substantial (though reduced amplitude) elongation response. Truncation of Component 0 by residual elongation response not yet suppressed at ISI = 0. 5 s by flashes bleaching 30% or more pigment also explains the smaller apparent magnitude of Component 0 for ISI = 0.5 s versus that for ISI = 1 s (Fig. 3D-H).

Given the previous attribution of cone outer segment elongation Component 2 measured in the single-flash paradigm to an opsin decay intermediate of bleached pigment (Pandiyan, Nguyen, et al. 2022b) (cf. SI, “1. Photosensitivity of ORG Elongation Response with the Single-Flash ORG paradigm and osmotic swelling”), it is reasonable to hypothesize that the positive-going phase of the ORGs obtained in response to stimuli with energy density exceeding 10^7^ photons μm^-2^ (and bleaching more than 50% of the pigment; Fig. 3B) is an early manifestation of Component 2, which is otherwise undetectable at early times in the single-flash paradigm due to the overwhelming elongation response. To test this hypothesis, we extended the kinetic scheme (Fig. 5H) for cone opsin decay to include two “swelling” states (C, D), with the latter following obligatorily from the “shrinkage” state (B) (Fig. 4I, Fig S3; SI, “Multi-state model of cone opsin bleaching and decay applied to Component 0 and Component 2 of the ORG”).

The key prediction of this hypothesis is that there are opposite-signed constants that linearly scale the occupancy of opsin states underlying both ORG negative-going shrinkage response and the slow elongation Component 2. Thus, in the absence of the fast elongation response (Component 1), the opsin multi-state-transition model predicts

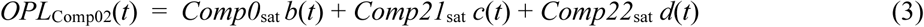

where the variables *b*(*t*), *c*(*t*), *d*(*t*) are the instantaneous occupancies of states B, C and D, *Comp0*_sat_ (negative), *Comp21*_sat_ (positive) and *Comp22*_sat_ (positive) are constant scaling coefficients, and the initial condition prior to the first flash in the dark-adapted eye is *P* = 1.0.

The multi-state opsin model was fitted to ORGs measured with ISI = 1 s (Fig. 6A-H, lower subpanels): here *Comp0*_sat_ *b*(*t*) is plotted as the dashed blue trace and the signed sum of Component 0 and Component 2 (Eq 3) as the unbroken blue trace. As some residual fast elongation response (Component 1) was also expected to be present (Fig. S1), a term *Comp1*(*t*) was added to Eq (3) for fitting the measured trace; the latter was constrained to have amplitude no greater than the residual predicted to be present 1 s after the initial flash. To address the status of elongation response, Δ*OPL*_Comp02_(*t*) (Eq 3) was subtracted from the “raw” data trace: the resultant “Comp02-corrected, pure Comp1” traces are presented in the upper of each pair of panels (Fig. 6C, 6E and 6F) to mean that the initial stimulus in these experiments effectively obliterated the fast elongation response (Component 1). The approximate correspondence of the estimated elongation response contributions at ISI = 1 s (red traces) in the lower panels of Fig. 6A and 6D and the Comp02-corrected traces (dark and light-blue traces in the corresponding upper panels) suggests that the substrate for elongation response was nearly completely depleted by 1 s after an initial 30% bleaching exposure (6.9×10^6^ photons μm^-2^).

**Figure 6.**
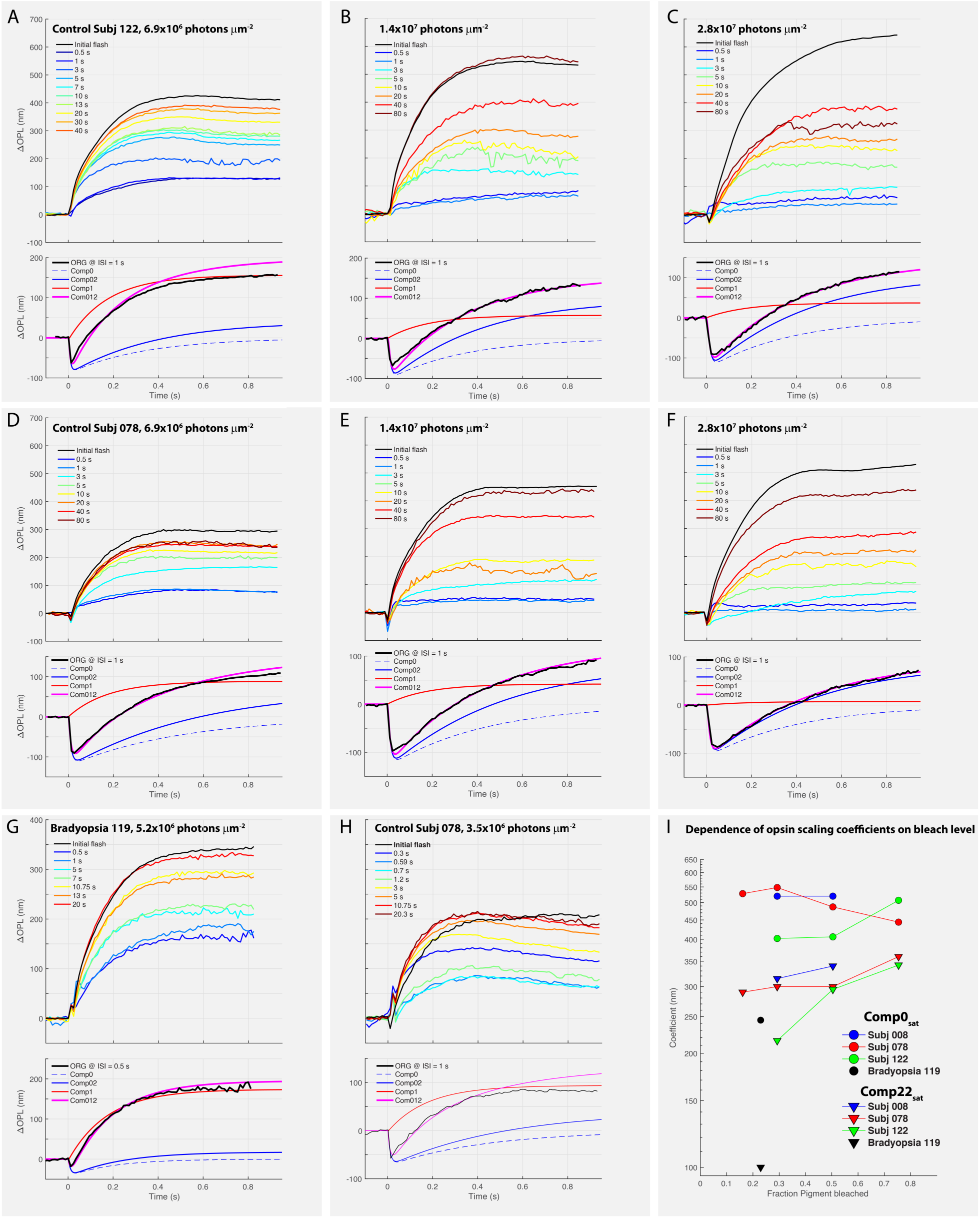
ORG Component 0 and early Component 2 can be understood as successive states of isomerized cone opsin, used to predict the time course of the initial response in the paired-flash paradigm, and to isolate the recovery of Elongation Response by subtraction. **A-H.** Each panel presents the complete set of traces from a paired-flash experiment (cf Fig. 2A); the subject and stimulus energy density of the stimulus used for the experiment are provided at the top of the panels. The lower subpanel in each case presents the ORG response obtained at ISI = 1 s (noisy black trace) and the component analysis the negative-going Component 0 is presented as a dashed blue line (“Comp0”), while the solid blue line presents the signed sum of Components 0 and 2 (Eq 3); the red trace presents Elongation Response (“Comp1”), and the magenta trace presents the sum of all components (“Comp012”) (cf legends on panels), which was fitted to the data trace. The upper panels in each case present the ORG data with the theoretical “Comp02” trace subtracted: this subtraction is rationalized by the hypothesis that Components 0 and 2 manifest successive states of the photopigment bleached by the 2^nd^ flash. Thus, the traces in the upper panel are predicted to represent solely the degree of recovery of Elongation Response at the ISIs given by the color bar on the panel. **I.** Dependence of the scaling parameters of the opsin state-transition model on the level of cone pigment bleaching for the bradyopsia patient (panel G), and for subjects for whom paired flash levels were performed with more than one flash strength.

The results of applying the multi-state opsin model are summarized in Fig. 6I. The scaling parameters *Comp0*_sat_ and *Comp22*_sat_ extracted from fitting the responses measured at ISI = 1 s are seen to be approximately independent of the bleach level produced by the initial stimuli but do exhibit systematic variation between subjects. The opsin-scaling parameters extracted from bradyopsia subject BO119’s single paired flash experiment (Fig. 6G) were notably smaller than those of the control subjects.

## DISCUSSION

### Normal RGS9 expression is required for deactivation of the human cone elongation response

Phototransduction in vertebrate rods and cones is realized by an amplified G-protein reaction cascade (Stryer 1986; Pugh and Lamb 1993; Arshavsky, Lamb, and Pugh 2002). The principal amplifiers of the cascade are photoisomerized opsin (R* in rods; P* in cones), which activates the cell’s heterotrimeric G-protein (G_t_α(GDP)βγ) by catalyzing GDP/GTP exchange, and cGMP phosphodiesterase (PDE6), activated by the binding of G_t_α-GTP. The cascade is deactivated by amplifier-specific reactions (reviewed in (Pugh and Lamb 2000)): R*/P* by C-terminal phosphorylation and binding of an arrestin (Xu et al. 1997; Chen et al. 1999; Lyubarsky et al. 2000; Nikonov et al. 2008) and the G_t_α(GTP)-PDE6 GAP complex by RGS9-1-catalyzed GTP hydrolysis (Lyubarsky et al. 2001; Krispel et al. 2006; Chen et al. 2000). Our investigation of light-driven elongation responses of cones in subjects deficient in R9AP, the anchor and transport protein for the regulator of G-protein signaling RGS9-1 (Baker et al. 2006; Nishiguchi et al. 2004), reveals that normal expression of RGS9 is required for normal deactivation of phototransduction in living human cones (Figs. 1, 2).

Light-driven cone elongation was previously hypothesized to be an osmotic swelling response (Zhang et al. 2017) to free phosphate (P_i_) produced by the G_tc_α-PDE-RGS9 complex GTPase reaction (Pandiyan, Nguyen, et al. 2022b). The responses of R9AP-bradyopsia subjects (Fig. 2A, B; Figs. S1, S2) reject this hypothesis. First, the P_i_ hypothesis predicts that the saturated amplitude of the elongation response would be reduced in the bradyopsia subjects, but the results show them to be normal (Fig. 2B, Fig. S2). Likewise, the P_i_ hypothesis predicts that the initial velocity of the elongation response for any given bleaching stimulus should be reduced in the bradyopsia subjects, but again, these were found to be normal. Nonetheless for its rejection, testing the P_i_ hypothesis in the bradyopsia subjects has proven to be highly informative in understanding the molecular mechanisms of the cone elongation response.

### Resting heterotrimeric G-protein is a substrate for the cone elongation response

Results presented here and previously situate the molecular substrate of the elongation response subsequent to P* activation of G_tc_, and before or at the RGS9-catalyzed GTPase deactivation of the PDE6 GAP complex in the phototransduction cascade, as follows. First, the rod homologue of the cone elongation response requires transducin, G_t_α (Zhang et al. 2017) and exhibits a qualitatively similar biphasic response profile consisting of a rapid shrinkage followed by an elongation (Li et al. 2026). Second, the elongation response is amplified relative to, and more photosensitive than bleaching (Zhang et al. 2017; Pandiyan, Nguyen, et al. 2022b)(Fig. 3, red curve), as expected for reactions of the G-protein cascade. Third, a substrate necessary for the elongation response is depleted in a bleach level-dependent manner (Fig. 3), consistent with the established evidence that the rate and quantity of activated G-protein increase with bleach (P*). Fourth, the similar amplitude and kinetics of cone elongation in bradyopsia subjects and controls (Fig. 2B) reveals that the amplified phototransduction product driving elongation is generated in the cascade sequence before completion of RGS9-catalyzed GTP hydrolysis. Fifth, the recovery of the elongation response amplitude after strong bleaches with a time course similar to that of pigment regeneration (Fig. 3I, K) (τ ∼ 2 min) (Hollins and Alpern 1973) implies that a long-lived intermediate species of bleached cone opsin, before it is quenched by regeneration, chronically depletes the substrate of the elongation response. In rods, persistent activation of G-protein (“*dark light*”) by bleached rhodopsin has long been established, with final recovery requiring regeneration by 11-*cis* retinal (reviewed in (Lamb and Pugh 2004)). In summary, the elongation response requires an amplified, P*- generated product that participates in the RGS9-catalyzed GTP hydrolysis reaction, identifying the resting heterotrimeric G-protein as the most plausible candidate.

### Recovery of the elongation response reveals remarkably fast cycling of G-protein in cones

The G-protein cycle in cones comprises at least seven distinct reactions beginning with the complex of P* and holo-G_tc_ and ending with the recombination of G_tc_α_2_(GDP) released by the RGS9- catalyzed GTP hydrolysis with Gβ_3_γ_8_. The cone dim-flash electrical response, like the rod single-photon response, has been long understood as key to the kinetics of the phototransduction reactions, as it reveals the molecular activation and deactivation processes operating in their linear response regime. Extending this framework to the elongation response under dim-flash conditions (Fig. 4) reveals an upper bound of ∼2 seconds for completion of the full G-protein cycle, a timescale consistent with the rapid G-protein cycling characteristic of phototransduction and approaching the fastest G-protein signaling cycles measured *in vivo* (Arshavsky, Lamb, and Pugh 2002; Ross and Wilkie 2000).

### Plausible biophysical mechanisms of the human cone elongation response

Though the evidence that resting heterotrimeric G-protein serves an essential substrate for the elongation response is strong, the specific biophysical mechanism by which elongation is produced remains unresolved. The following three hypotheses appear to be currently tenable.

#### Hypothesis 1: partitioning of water into the P*-G_tc_ complex

A biophysical basis for the elongation response that maintains a role for G-protein signaling is the water-stabilized volume increase of the complex of photoisomerized cone opsin (P*\**) and the heterotrimeric G-protein complex (*G*_tc_-GDP) (Chawla et al. 2021). This complex must exist long enough for the guanine nucleotide binding cavity in *G*_tc_α_2_ to open and release GDP and GTP to enter and bind. The molecular unit of the water activity increase should include not only the binding pocket of *G*_tc_α_2_ for guanine nucleotide and the “vestibule” of the GDP/GTP exchange, but also the volumetric change of water in P* (Chawla et al. 2021). The rationale for including the P* component of the P*-G_tc_ activation complex in the water unit is that this complex exists as a molecular unit (Gao et al. 2019; Yao, Fay, and Farrens 2025), G_t_α binding stabilizes Metarhodopsin II (Emeis et al. 1982; Hofmann et al. 2009), and the formation of the complex increases water imbibition by Metarhodopsin II (Chawla et al. 2021).

#### Hypothesis 2: cone outer segment (COS) refractive index change

The binding of R* to rod G_t_ causes an increase in infrared light scattering from rod disc membranes – the so-called “binding signal” (Kuhn et al. 1981) – attributable to a change in the refractive index at the disc membrane surface (reviewed in (Arshavsky, Lamb, and Pugh 2002)). An increased condensation of protein at the disc membrane surface can alter the bulk refractive index of the outer segment (Corless and Kaplan 1979; Kaplan 1982; Kaplan and Liebman 1977). Such a mechanism would be expected to have kinetics similar to that of the “sponge-like” increases in water activity accompanying formation of the P*-G_tc_ complex.

#### Hypothesis 3: *G*_t_α-GTP triggered *PDE6 elongation*

The cryoEM structure of PDE6α/β with two G_t_α subunits bound is observed to be elongated by ∼ 4 nm relative to the apo-structure (7JSN) (Gao et al. 2020), consistent with a recent study suggesting a link between PDE6 activation and outer segment elongation (Tomczewski et al. 2025). Complete activation of PDE6 in the outer segment could therefore underlie the saturated 460 nm increase in cone outer segment length, corresponding to ∼ 0.5 nm per interdiscal space for ∼800 discs (Table 1). Challenges faced by this hypothesis include (i) whether PDE6c spans the intradiscal gap in cone outer segments and, if not, by how much; (ii) whether the elongation measured in the cryoEM structure occurs *in vivo* and (iii) whether, given its relative paucity, PDE6 elongation can generate sufficient force in the presence of interdiscal elasticity (Haeri, Knox, and Ahmadi 2013) to account for both the maximal cone elongation and its photosensitivity (Fig. S1).

**Table 1.** Parameters of the Multi-state Opsin Model Applied to Paired-Flash.

| *Subject* | *Test stimulus*<br>(photons $\mu\text{m}^{-2}$) | *1<sup>st</sup> fraction bleached* | *2<sup>nd</sup> fraction bleached* | *Comp0<sub>sat</sub>*<br>(nm) | *Comp22<sub>sat</sub>*<br>(nm) | *Comp1* (nm)<br>(ISI = 1 s) | $k_0$ (s<sup>-1</sup>) | $k_1$ (s<sup>-1</sup>) | $k_2$ (s<sup>-1</sup>) | $k_3$ (s<sup>-1</sup>) | *Figures* |
| --- | --- | --- | --- | --- | --- | --- | --- | --- | --- | --- | --- |
| Bradyopsia 119 | $5.2 \times 10^6$ | 0.23 | 0.18 | -233 | 100 | 177 | 74 | 100 | 5.0 | 5.5 | 6G, 6I |
| Control 008 | $6.9 \times 10^6$ | 0.29 | 0.21 | -536 | 300 | 128 | 88 | 96 | 1.8 | 2.7 | 6I |
| Control 008 | $1.4 \times 10^7$ | 0.50 | 0.29 | -520 | 340 | 59 | 110 | 96 | 1.8 | 2.8 | 6I |
| Control 078 | $3.5 \times 10^6$ | 0.16 | 0.14 | -528 | 290 | 99 | 88 | 90 | 2.3 | 3.0 | 6H, 6I |
| Control 078 | $6.9 \times 10^6$ | 0.29 | 0.21 | -548 | 300 | 90 | 88 | 95 | 2.0 | 4.0 | 6D, 6I |
| Control 078 | $1.4 \times 10^7$ | 0.50 | 0.27 | -488 | 300 | 50 | 98 | 95 | 2.3 | 4.4 | 6E, 6I |
| Control 078* | $2.8 \times 10^7$ | 0.75 | 0.24 | -444 | 360 | 7.5 | 203 | 95 | 2.5 | 4.0 | 6F, 6I |
| Control 078* | $2.8 \times 10^7$ | 0.75 | varied 2<sup>nd</sup> fl. | -426 | — | — | 220 | 93 | — | — | 5H |
| Control 122 | $6.9 \times 10^6$ | 0.29 | 0.21 | -402 | 180 | 135 | 97 | 150 | 3.0 | 5.5 | 6A, 6I |
| Control 122 | $1.4 \times 10^7$ | 0.50 | 0.29 | -406 | 295 | 57 | 122 | 110 | 3.0 | 4.4 | 6B, 6I |
| Control 122* | $2.8 \times 10^7$ | 0.75 | 0.009 | -507 | 360 | 55 | 201 | 95 | 3.5 | 4.4 | 6C, 6I |
| Average | — | — | — | -480 | 316 | — | — | 102 | — | — |  |

### Confirmation of the opsin basis of Components 0 and 2 of the optoretinogram (ORG)

The cone ORG comprises not only the fast elongation response (“Component 1”), but also two other kinetically distinct features, an initial rapid optical path length decrease (“Component 0”) and a slower optical path length increase (“Component 2”) (Pandiyan, Nguyen, et al. 2022a). Results presented here confirm that species of photoactivated cone opsin underlie both ORG Components 0 and 2 and extend their kinetic characterization (Figs. 5, 6). The paired-flash paradigm, by suppressing the fast, truncating elongation response, reveals that the saturated magnitude of negative-going Component 0 exceeds 100 nm, and that the latter’s magnitude is reduced in a bleach level-dependent manner (Fig. 5). The paired-flash results also give a clearer picture of Component 2’s kinetics and indeed suggest that the opsin state that underlies Component 0 transitions directly into a distinct state underlying Component 2 (Fig. 6). Classically, opsin states have been identified by spectroscopic signatures and more recently structural models, but Component 0 and Component 2 could manifest previously unrecognized states resulting from changes in opsin hydration.

### The optoretinogram (ORG): a clinically translatable assay of photoreceptor function and G-protein signaling

The ORG enables sensitive, non-invasive characterization of cone function in the macula, the retinal region responsible for high-acuity, daylight vision and a substantial fraction of visual cortical input. As such, it provides a quantitative means to monitor the functional status of cone photoreceptors in both healthy and diseased retina (Liu et al. 2025; Wendel et al. 2024; Lassoued et al. 2021; Gaffney, Connor, and Cooper 2024; Xu et al. 2024) and has potential for application in progressive conditions such as age-related macular degeneration and during therapeutic intervention. By establishing the molecular basis of the ORG, the present work provides a framework to link functional measurements in living human photoreceptors to the underlying mechanisms of G-protein signaling in native and diseased conditions. More broadly, ORG recordings from human subjects carrying transduction-protein mutations provide an avenue, analogous to classical electrophysiological recordings in genetically modified animal models, for identifying and quantitatively characterizing specific elements of the phototransduction cascade *in vivo*.

## MATERIALS AND METHODS

### Subjects

Four control subjects (two males and two females, aged 31–43) with no known retinal disease, along with two female siblings, each carrying a homozygous in-frame deletion (c.94_102del; p.Asp32_Gln34del) in the *R9AP* gene associated with bradyopsia (Figure 1), were enrolled in this study. Pupillary dilation was achieved using 1% tropicamide ophthalmic solution (Akorn Inc.) and/or 1% phenylephrine. The study was approved by the University of Washington Institutional Review Board and conducted in accordance with the tenets of the Declaration of Helsinki. Written informed consent was obtained from all participants after explaining the study procedures and potential risks, but prior to their participation.

### Immunochemistry

#### Constructs

RGS9-1 (GenBank: NM_001165933) in pcDNA3.1(+) were synthesized by GenScript (Masuho et al. 2020). Gβ5L (GenBank: NM_016194) in pcDNA3.1(+) were purchased from cDNA Resource Center (https://www.cdna.org) (Masuho et al. 2020). C terminal FLAG-tagged R9AP-wt in pcDNA3.1(+) (GenBank: NM_207391.3, OHu19327D) were purchased by GenScript. C terminal FLAG-tagged R9AP-del (94_102del) in pcDNA3.1(+) were synthesized by GenScript.

#### Antibodies

Rabbit anti-Gb5 (ATDG) antibody was the generous gift from Dr. William Simonds [National Institute of Diabetes and Digestive and Kidney Diseases/National Institutes of Health (NIH), Bethesda, MD]; rabbit anti-RGS9-1 antibody was the generous gift from Dr. Theodore Wensel, [Biochemistry and Molecular Pharmacology, Baylor College of Medicine, Houston, TX]. Commercial antibodies were as follows: Monoclonal ANTI-FLAG® M2-Peroxidase (HRP) antibody (A8592, Sigma-Aldrich); mouse anti-b-actin (AC-15, Sigma-Aldrich).

#### Transfection

Cells were seeded in a 6-well plate at a density of 2×10⁶ cells per well for transfection. After 3 hours, expression constructs (2.5 mg total per well) were transfected into the cells using Lipofectamine LTX reagents. An empty vector (pcDNA3.1(+)) was included to normalize the total amount of transfected DNA.

#### Immunoprecipitation and Western blotting

Transfected HEK293T cells from one well of a 6-well plate were lysed with 500 μl of PBS supplemented with 150 mM NaCl, 1% Triton X-100, and Complete protease inhibitors (Roche) and homogenized by sonication. The lysate was centrifuged at 20,800 × g for 15 minutes at 4°C. The resulting supernatant was incubated with 20 μl of anti-FLAG affinity resin (L00432, GenScript) on a rocker at 4°C for 1 hour. After three washes with lysis buffer, proteins were eluted from the beads using 50 μl of SDS sample buffer containing 8 M urea, incubated for 10 minutes at 42°C. The eluted proteins were analyzed via SDS-PAGE, followed by Western blotting using specific antibodies and HRP conjugated secondary antibodies and an ECL (Kindle Biosciences) detection system. The following dilutions of the antibodies were used: rabbit anti-RGS9-1, 1:5000; rabbit anti-Gb5 (ATDG), 1:2500; anti-FLAG® M2-Peroxidase (HRP), 1:5000; mouse anti-β-actin, 1:5000. Signals were captured on KwikQuant Imager (Kindle Biosciences). For quantification, band intensities were determined by using NIH ImageJ software. The integrated intensity of β-actin was used to normalize data for lysate loading, while the integrated intensity of R9AP in the elution was used to normalize data for immunoprecipitation.

### Long-read genome sequencing

The two bradyopsia study subjects were consented for targeted long-read genome sequencing under an approved protocol by the IRB at the University of Washington, Seattle, WA. Written informed consent was obtained for a venipuncture to obtain 2 milliliters of blood. Genomic DNA was isolated for long-read library preparation and targeted long-read inherited retinal disease panel sequencing with adaptive sampling for targeted enrichment and variant level analysis using previously published methods (Nakamichi et al. 2024). Previous studies had used primer-guided PCR amplification of exons of *R9AP* (14), whereas in this case targeted long-read sequencing of the RGS9 and R9AP locus and surrounding region was performed. It was concluded that the only alteration in the subjects was the previously identified homozygous in-frame deletion mutation in R9AP, thereby excluding any additional genetic or epigenetic alterations contributing to their bradyopsia.

### Adaptive optics scanning light ophthalmoscopy

Images of the cone mosaic were obtained using a previously described confocal adaptive optics scanning light ophthalmoscope (AOSLO)(Dubra and Sulai 2011). The imaging source was a superluminescent diode (SLD, Superlum, Carrigtwohill, Ireland) with a center wavelength of 775 nm, while an 850 nm SLD was used as the wavefront sensing source. Using two square FOVs (1.0° x 1.0° and 1.5° x 1.5°), image sequences of 150 frames each were acquired at multiple retinal locations. Each image sequence was processed as described by (Salmon et al. 2017). The processed images were then montaged using an automated algorithm(Chen et al. 2016), and the resultant montage was manually reviewed and adjusted in Photoshop (Adobe Creative Suite, San Jose, CA). The location of highest cone density (tightest cone spacing) was marked as the (0,0) center, and regions of interest (ROIs) were extracted from the montage using Mosaic Analytics (Translational Imaging Innovations, Hickory, NC, USA). The size of each ROI was sized based on the known relationship between cone density and retinal eccentricity, such that each ROI would be expected to contain about 100 cells. Cones in each image were then semi-automatically identified using Mosaic Analytics (Translational Imaging Innovations, Hickory, NC, USA), and cone density was computed for each ROI as the number of cones whose Voronoi area vertices were contained within the ROI, divided by the summed area of these “bound” Voronoi areas (Cooper et al. 2016).

### Optoretinography – optical imaging system

A previously described coarse-scale optoretinography (CoORG) system was used to measure cone ORGs (Jiang et al. 2022). The CoORG setup includes three modules: a spectral-domain optical coherence tomography (OCT) system, a line-scan ophthalmoscope (LSO), and a stimulus delivery module for retinal illumination. A superluminescent diode (λ = 840 nm, Δλ = 50 nm; MS-840-B-I-20, Superlum, Ireland) provided the light source for both the OCT and LSO. The 3D retinal volumes were acquired in Fourier domain using a spectrometer comprising a 1200 line-pairs/mm diffraction grating, an imaging lens, and a high-speed 2D CMOS camera (Photron, FASTCAM NOVA S6). The LSO used a line-scan camera (Basler, Sprint spL2048-70km) to capture *en face* retinal images and assisted in optimizing focus. A 528 ± 5 nm LED delivered the visual stimulus in Maxwellian view for the ORG experiments.

### Stimulus calibration and photon density calculation

Light stimulus calibration and photon density calculation followed previously published protocols (Jiang et al. 2022; Pandiyan, Schleufer, et al. 2022). The LED spectrum and output power were measured and converted to photon flux. Corrections for ocular media absorption—including lens and macular pigment—were applied using established values from literature. The corrected photon flux was then normalized by the stimulus area in angular units and converted to retinal photon density (photons/µm²) using a retinal magnification factor that was adjusted based on each subject’s axial length and dilated pupil size.

### ORG Protocol

Subjects were initially imaged using the line-scan ophthalmoscope (LSO) modality to identify the appropriate region of interest (ROI) and achieve optimal focus in the outer retina. A reference LSO video was recorded, after which the system was switched to the line-scan optical coherence tomography (OCT) modality. Both the LSO and OCT imaging covered a 5° × 5° field of view (FOV), centered at 4° temporal eccentricity. Before each ORG experiment, subjects were dark-adapted for 2 minutes. OCT volumes were acquired using two stimulus protocols described below: the single-flash and paired-flash paradigms. The single-flash paradigm was designed to probe the activation dynamics of ORG components. In contrast, the paired-flash paradigm was used to study the recovery of the ORG, and to further characterize Component 0 by suppressing Elongation Response.

#### Single Flash ORG Paradigm

OCT volumes were acquired under two conditions differentiated by the length of recording. In the short-duration protocol, volumes were acquired at 16,000 B-scans per second, with each volume containing 500 B-scans. Ten pre-stimulus volumes were recorded, followed by 40 post-stimulus volumes. For long-duration recordings, the acquisition rate was reduced to 12,000 B-scans per second, with each volume comprising 400 B-scans. The protocol included 10 pre-stimulus volumes followed by 210 post-stimulus volumes acquired over a 13-second period. Immediately after the stimulus, volumes were continuously recorded for 2.3 seconds, after which acquisitions resumed at 1.5- second intervals, with a 3-second pause between subsequent recordings. This acquisition strategy allowed for extended temporal tracking of ORG responses while remaining within the bandwidth and memory buffer size of the camera.

#### Paired-flash ORG Paradigm

Two stimuli—referred to as the “test” and “probe” flashes—were delivered with a variable inter-stimulus interval (ISI) as shown in Figure 3A. OCT volumes were acquired at a rate of 16,000 B-scans per second, with each volume consisting of 160 B-scans, yielding a volume rate of approximately 100 Hz over a 3° × 5° FOV. A total of 210 volumes were acquired, including 10 pre-stimulus volumes and 200 post-stimulus volumes after the combined test and probe flashes. For measuring the recovery of ORG, the ISI was varied between 0.5 and 80 seconds, while the intensity of both test and probe flashes was held constant. For specific characterization of Component 0, the ISI was fixed at 1 second, the duration of the test flash was either 4 or 16 ms, and the probe flash duration was systematically varied between 1- 50 ms. The change in duration was used to adjust retinal photon density and bleach strength, given the limit on maximum power emanating from the stimulus LED source.

### ORG Data processing

Image processing followed established methods as described in (Jiang et al. 2022). Briefly, raw data were reconstructed in k-space, and a fast Fourier transform (FFT) was applied to generate B-scan stacks for each OCT volume. Retinal layer segmentation was performed using open-source software. Trials with identical stimulus conditions were co-registered using a strip-based registration algorithm using the *en face* image at the inner segment/outer segment (ISOS) junction. From the registered volumes, *en face* images were extracted at the ISOS and cone outer segment tips (COST) layers. Phase differences between these two layers were computed, and ORGs were derived by averaging the phase differences over the region of interest. These phase differences were then converted to changes in optical path length (ΔOPL).

For analysis of ORG response recovery dynamics and characterizing Component 0, the following parameters were extracted: initial slope, maximum amplitude, and Component 0 amplitude. ORG response recovery was defined as the amount of time it takes for the ORG amplitude and initial slope to reach the same level as the initial test flash. The Component 0 amplitude was quantified as the local minimum in ΔOPL relative to the pre-stimulus volume.

## ACKNOWLEDGEMENTS

Supported by National Eye Institute grants U01EY025501, R01EY029710, R01EY018139, R01EY017607, P30EY001730, Research to Prevent Blindness, Gene & Ruth Posner Foundation, George and Martina Kren Endowed KEMI Chair in Vision Research, Kren Engineering in Medicine Initiative, SPIE Franz-Hillenkamp Postdoctoral Fellowship.

---

**Table S1.** Quantitative Characteristics of ORG Elongation.

| *Subject* | $L_{\text{COS}}$ ($\mu\text{m}$) | $\Delta OPL_{\text{sat}}$ (nm) | $\Delta L_{\text{sat}}$ (nm) | $\Delta V$ ($\mu\text{m}^3$) | $Q_e$ (photons $\mu\text{m}^{-2}$) | *Figures* |
| --- | --- | --- | --- | --- | --- | --- |
| Control 008 | 21.6 | 590 | 420 | 1.0 | $6.6 \times 10^6$ | 3D, 3E; 4F-4J; S1 |
| Control 078 | 20.5 | 525 | 372 | 0.9 | $6.6 \times 10^6$ | 2A, 2B; 4A-4E; 5A-5E; 6D-6H; S1 |
| Control 112 | 22.0 | 700 | 496 | 1.2 | $6.6 \times 10^6$ | 2A, 2B; S1 |
| Control 122 | 21.8 | 720 | 511 | 1.2 | $6.6 \times 10^6$ | 2A, 2B; 3F-3H; 5C, F; 6A-6C; S1 |
| Bradyopsia 119 | 23.3 | 650 | 461 | 1.1 | $6.6 \times 10^6$ | 2A, 2B; S1, S2 |
| Bradyopsia 120 | 22.0 | >450 | NA | NA | NA | 2A; S2 |

## SUPPLEMENTARY INFORMATION

## 1. Photosensitivity of ORG Elongation Response with the Single-Flash ORG paradigm and osmotic swelling

ORGs of the subjects of the study were obtained in the single-flash paradigm previously used to define Elongation Response (Pandiyan, Nguyen, et al. 2022b) (Fig. S1). Rising exponential functions with a fixed time constant were fitted to the traces as in (Pandiyan, Nguyen, et al. 2022b), and the amplitudes plotted as a function of the stimulus energy density and fitted with a saturating exponential (Fig. S1D). Elongation response of all observers had a common photosensitivity at this eccentricity, 1/*Q*_e,C1_=1.5×10^−7^ μm^2^, 3-fold greater than the photosensitivity of bleaching, 1/*Q*_e_ =5×10^−8^ μm^2^ (Fig. 3B). The saturated amplitude of the exponentials also provides an estimate of the osmolarity change underlying the responses, predicted on the assumption that these represent osmotic equilibria. Applying Van’t Hoffs law of osmotic equilibrium and the assumption that the COS outer segment cytosol is 50% of the envelope volume, one can readily obtain the relation Δ*C* = 300 × [Δ*L*/(*L*_cos_/2)], where Δ*C* (units: mOsM) is the incremental change in COS osmolarity, 300 (mOsM) is the osmolarity of blood plasma, *L*_cos_ (nm) is the length of the COS in the dark adapted state and Δ*L* (nm) is the length change measured on the plateau. Rearranging the terms, one obtains the scaling constant Δ*C/*Δ*L* = 0.027 mOsM nm^-1^. Apply this scaling factor to the average saturated Component 1 length increase adjusted for the refractive index of the outer segment (n = 1.41), one obtains, Δ*C* = (650 nm /1.41) × 0.027 mOsM nm^-1^ = 12.4 mOsM, a 4% increase in osmolarity.

**Figure S1.**
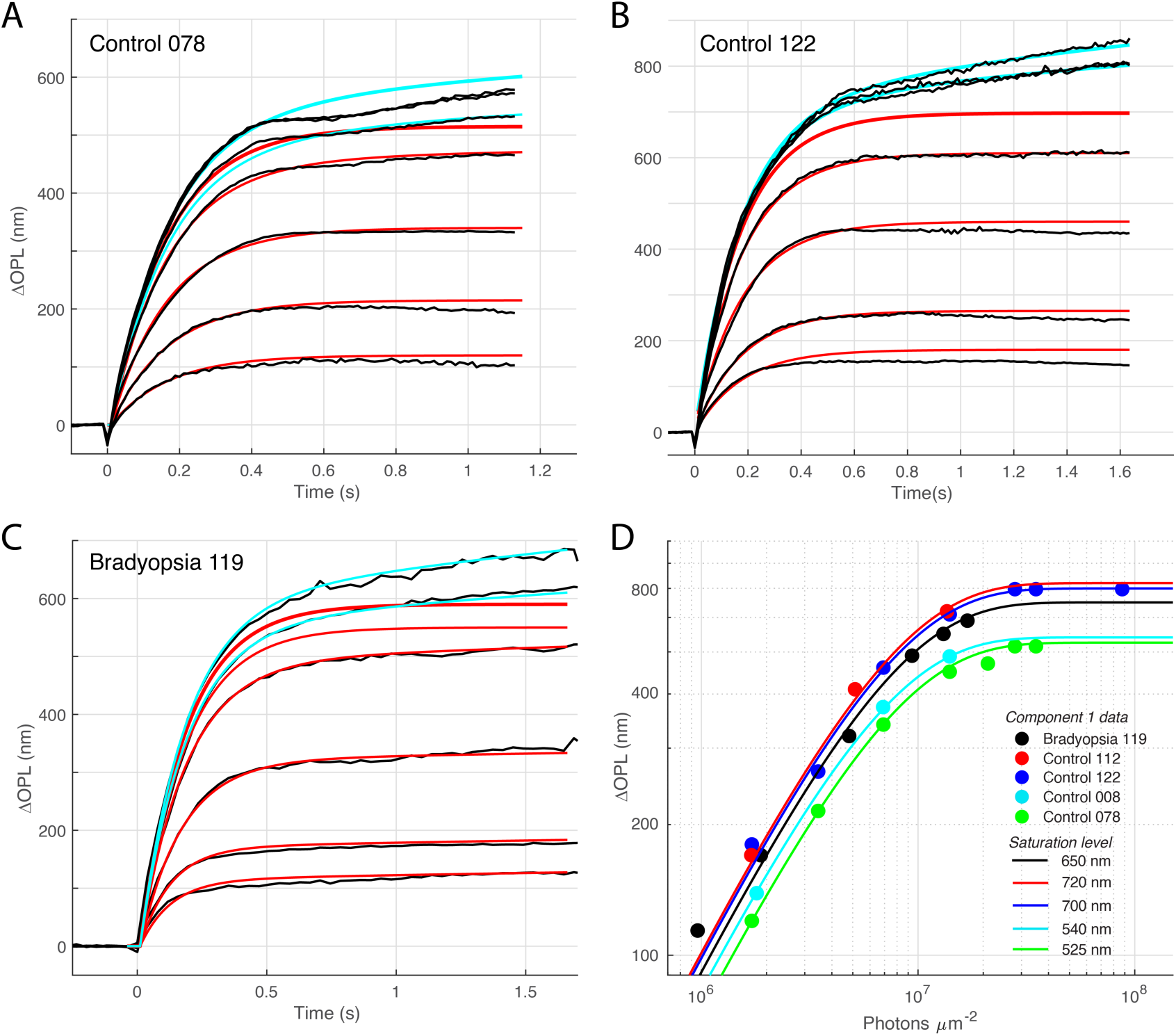
Single-flash ORG response families of the principal subjects of the study. **A-C.** ORG data (black traces) were fitted with rising exponential curves, as previously (Pandiyan, Nguyen, et al. 2022b): the lower amplitude traces were well described by a single rising exponential curve, which had the time constants 175 ms (Subj 122), 145 ms (Subj 078) and 156 ms (Bradyopsia 119), and the same exponential function (Elongation Response) was fitted to the three uppermost curves along with a more slowly rising component (Component 2), with time constants (0.5 s, 0.8 s and 5 s, respectively); the cyan traces are the sum of the two components. The red curves describe Elongation Response, and the cyan curves plot the sum of Elongation Response and Component 2 (as described in ref (Pandiyan, Nguyen, et al. 2022b)). The topmost thickened red trace shows the saturated Elongation Response obtained from the fitting. (Component 2 is not shown separately, as more intense stimuli are needed to generate it maximal amplitude). **D.** Response amplitude vs. intensity data from panels A-C (symbols) and exponential saturation functions *R* = *R*_max_(1-exp(-*Q*/*Q*_e,C1_)] fitted to the Elongation Response amplitude data (and to data of Subj 112, not shown). The saturation functions all have the same value of *Q*_e,C1_, 6.6×10^6^ photons μm^-2^. As cone pigment bleaching measured directly by “Adaptive-optics, serial-scanning single-cell reflectometry” (AO-SSSCR) (Pandiyan, Nguyen, et al. 2022b) was found have a photosensitivity 1/ *Q*_e,bleach_ = 1/ (2×10^7^ photons μm^-2^) (Fig. 3B), Elongation Response is 3-fold more photosensitive than bleaching at 4.5 deg temporal eccentricity in these subjects. (There were insufficient data – cf. Fig. S2 – from the second bradyopsia subject for a response vs. intensity analysis of Elongation Response.)

## 2. Complete ORG results from bradyopsia subjects

**Figure S2.**
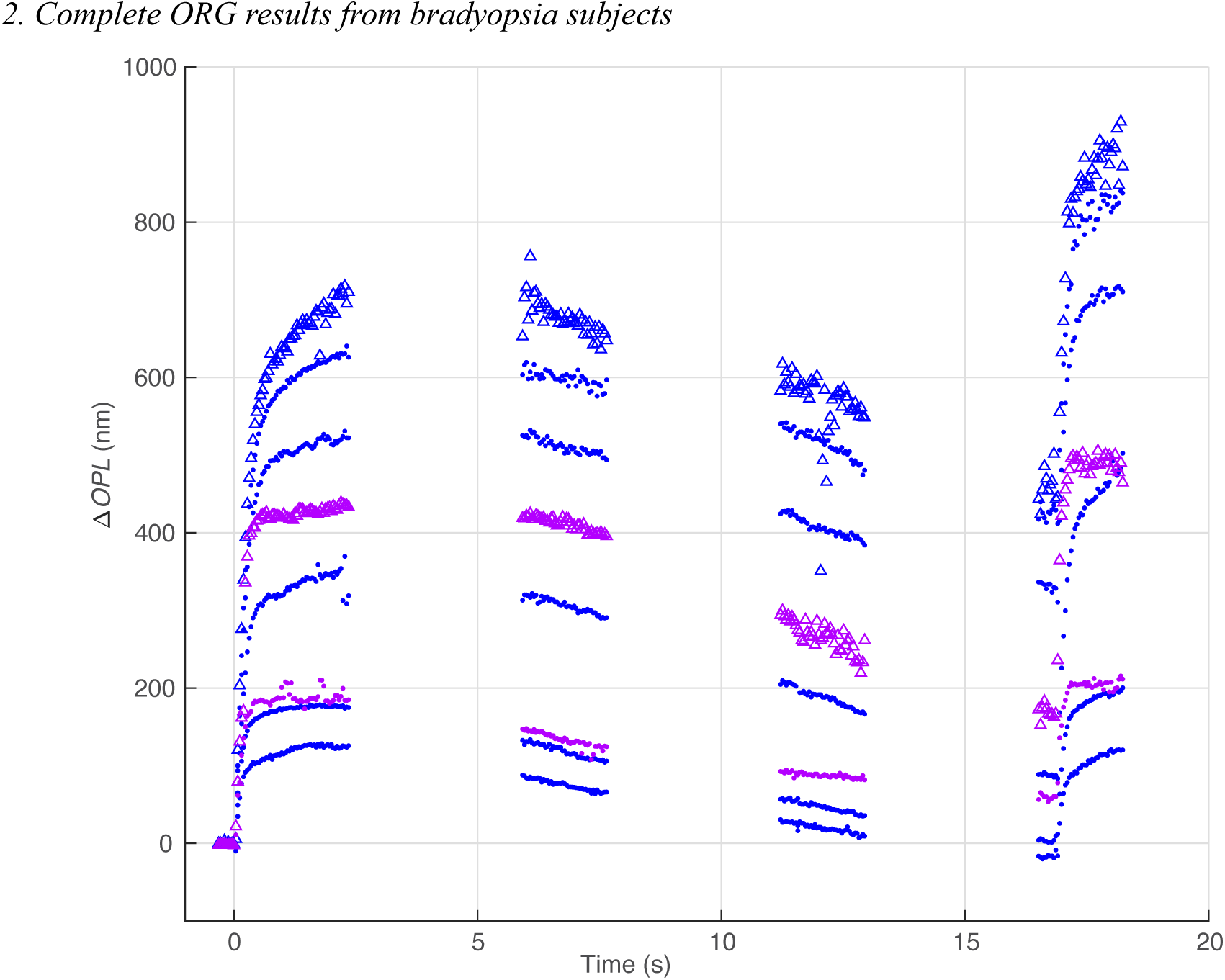
Single-flash long-term ORGs of the two bradyopsia subjects: BO119 (blue symbols); BO120 (purple symbols). The lowermost three traces are also presented in Fig. 2A, B, and the response vs. intensity (photosensitivity) analysis is presented in Fig. S1. (Open symbols used to help distinguish overlapping records.) The lowermost three traces are presented and analyzed in Fig. 1A, B for their recovery kinetics.

## 3. Multi-state model of cone opsin bleaching and decay applied to Component 0 and Component 2 of the ORG

Photoisomerized opsins have long been known to undergo successive “thermal decay” through several distinct states. Classically, these states were identified spectroscopically – thus, photoisomerized rhodopsin transitions through lumi-rhodopsin, Metarhodopsin I ←→ Metarhodopsin II ←→ Metarhodopsin III (the double arrows indicate equilibria); the isomerized chromophore all-*trans* retinal remains covalently linked to its Schiff-base in the lumi- and meta- states, but ultimately must be hydrolyzed, exit the binding pocket and be replaced by fresh 11-*cis* retinal to reform the functional pigment. Inspired by the work of Chawla et al. (Chawla et al. 2021), we previously hypothesized that ORG Component 2 could represent outer segment swelling caused by a water-imbibed state of cone opsin. Here, we adopt a more general strategy to accommodate the initial ORG Component 0. Specifically, we suppose that isomerized cone pigment molecules transition successively from their regenerated pigment state P through 4 successive states A, B, C, D as follows:

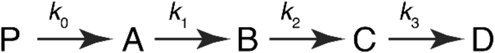

This scheme can readily be implemented for a homogenously illuminated outer segment as a series of first-order differential equations and solved analytically. The rate equation for the first step is familiar in the visual pigment literature, but bears repeating because of a facet of the experimental protocol, whereby different degrees of bleaching were achieved by varying the duration of the light pulse. Specifically, we write the rate equation for bleaching as

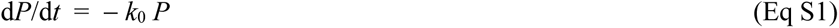

with *k*_0_ = *I*_0_ /*Q*_e_, where *I*_0_ (photons μm^-2^ s^-1^) is the power density of the light at the retinal surface and *Q*_e_ is a constant. Subject to the initial condition *P = P*_0_ = 1 for *t* ≤ 0, the solution to Eq S1 for a light pulse of duration Δ*T* s is

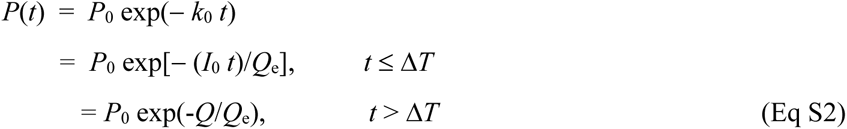

where *Q* = *I*_0_ Δ*T* is the energy density (photons μm^-2^) of the pulse. The importance of this exposition for the present paper is two-fold. First, depending on *I*_0_ and *Q*_e_, exposures exceeding a certain time Δ*T\** produce negligible additional bleaching with increase duration, as the pigment will be effectively completely bleached by some duration. For example, for a 90% bleach, Δ*T\** = (*Q*_e_ /*I*_0_) log_e_(0.01) ∼ 26 ms for the dark-adapted photosensitivity. Exhaustive bleaching for the longer duration exposures explains the saturation behavior in Fig. 5H. A second reason for the importance of Eq S2 is that bleaching by the initial stimulus in the paired flash experiments affects the quantity of pigment bleached by the second flash, and also sets the initial conditions of the variables for the state occupancy at the second flash. One way to illustrate this is with a plot of the state variables for flashes of varied duration.

**Figure S3.**
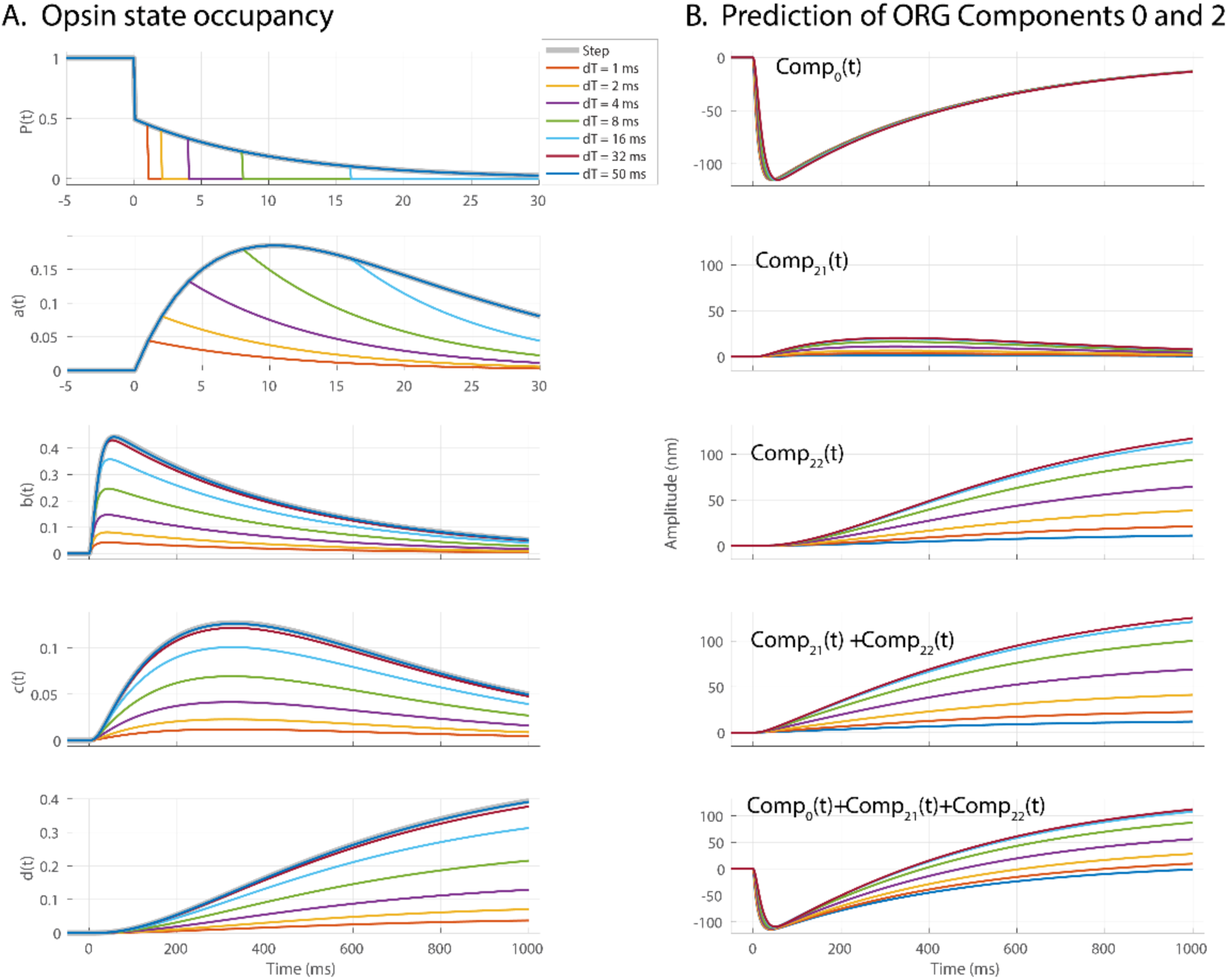
Multi-state cone opsin decay model and predictions of the unified Component 0 and Component 2. **A**. Occupancy of the states of the opsin bleaching and decay scheme: the variables in the 5 panels correspond to the 5 states in the transition schematic above. The topmost variable, *P*(*t*) is the fractional quantity of pigment present in the cone outer segment: bleaching commences at *t* = 0 in response to light whose power density is 1.75×10^9^ photon μm^-2^ s^-1^, and is terminated after different pulse durations dT varying from 1 to 50 ms. Note that the two upper panels (for the variables *P*(*t*) and *a*(*t*)) are presented on a 30 ms axis scale, but the remaining state variables are presented on 1 s scale, corresponding approximately to the ORG data. The rate parameters employed are those for the paired flash experiment for Control Subject 078, 5^th^ row of Table 1. **B**. Prediction of the time course of Components 0 and 2, as per Eq 1 of the text. The trace in the lowest panel corresponds to that used in the prediction of “Comp02” in Figure 6E.

## 4. Increase in the photosensitivity of bleaching by strong prior bleaches

As reported in the presentation of the results in Figure 5G and 5H, the photosensitivity of ORG Component 0 exceeds that of cone pigment bleaching at the same retinal locus by 2.5-fold: the increased photosensitivity is seen in Fig. 5G as the leftward shift between the unbroken black curve describing the ORG data and the dashed curve, which plots the fraction bleached by the stimuli whose energy density is given by the abscissa: this curve describing bleaching (*B*) is the complement of the curve in Fig. 3B (i.e., *B* = 1 – *P*) and is plotted in Fig. 5G (dashed line) scaled to the saturating magnitude of that describing Component 0. One mechanism expected to produce a *Q*_e_-shift of this sort was described by Pandiyan *et al*., (Pandiyan, Nguyen, et al. 2022b), Fig. S3 in reference), and arises from the change in pigment axial density attendant bleaching (Fig. S4). The *Q*_e_-shift predicted by this mechanism for the experiment of Fig. 5 contributes to the observed shift, but cannot fully explain the ORG Component 0 *Q*_e_-shift of Fig. 5G, as the predicted shift is unlikely to exceed 1.5-fold (panel C, red symbol). Another mechanism by which bleaching could shift the *Q*_e_ for bleaching is a change in the coupling efficiency of waveguiding between inner and outer segment, for which there is some evidence. Thus, Miller (Miller 1972) apparently found that the Stiles-Crawford parabola broadens during bleaching: such broadening suggests light capture by the cone waveguide from a greater fraction of the pupil area after substantial bleaching as compared to capture in the dark-adapted state.

**Figure S4.**
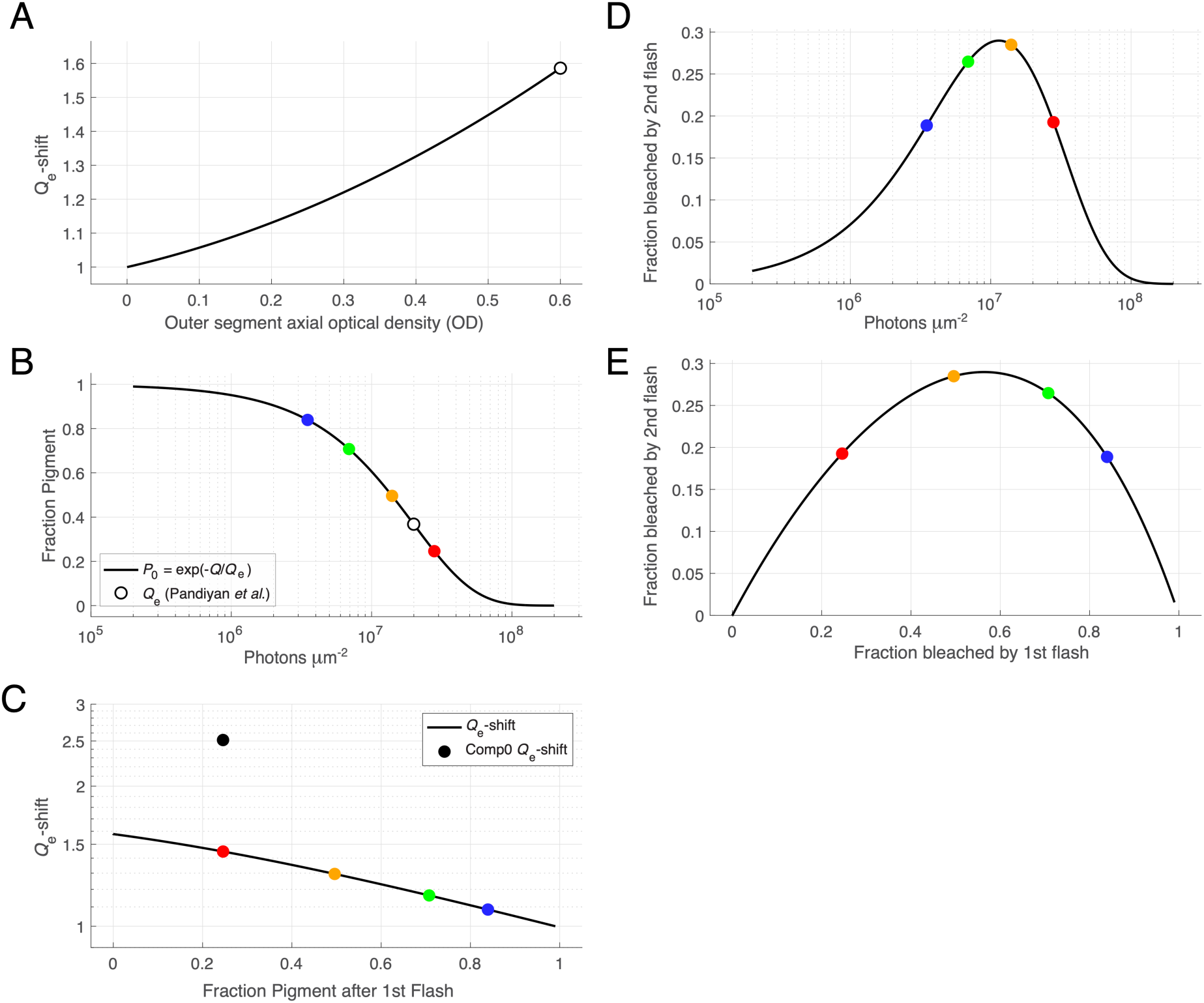
Decreased cone pigment axial density caused by bleaching generates a *Q*_e_-shift for subsequent bleaching exposures. **A.** The *Q*_e_-shift between cones whose outer segments have different total axial optical density, plotted for cones with a maximal density of 0.6 OD units (replotted from (Pandiyan, Nguyen, et al. 2022b), Fig. S3D; cf. also Miller (Miller 1972)) **B.** Curve describing the energy density dependence of cone pigment at 4.5 deg temporal eccentricity as measured by Pandiyan *et al*. (Fig S2 in (Pandiyan, Nguyen, et al. 2022b)); the colored symbols plot the calculated bleach fractions for the energy densities of the stimuli used in the paired-flash experiments, with the same color coding (cf. Fig. 3K). **C.** Replotting the curve of panel A as a function of the fraction pigment bleached by an initial flash in the paired-flash experiments: bleaching by an initial stimulus delivering 2.8×10^7^ photons μm^-2^ (red symbol) is predicted to shift the *Q*_e_ lower for bleaching by a second flash ∼1.5-fold, substantial but less than the observed 2.5-fold shift of Component 0 (filled black symbol). **D, E.** Calculations of the fraction of pigment bleached by the second flash in a paired flash experiment when the first flash has the same energy density, and the *Q*_e_ is shifted 2.5-fold, plotted as a function of the energy density (D) or the initial bleach fraction. (Note that bleach fraction is calculated in reference to the total quantity of pigment in the dark-adapted cone, taken to be unity; cf notes to Table S1).

